# Active Learning of the Conformational Ensemble of Proteins using Maximum Entropy VAMPNets

**DOI:** 10.1101/2023.01.12.523801

**Authors:** Diego E. Kleiman, Diwakar Shukla

## Abstract

Rapid computational exploration of the free energy landscape of biological molecules remains an active area of research due to the difficulty of sampling rare state transitions in Molecular Dynamics (MD) simulations. In recent years, an increasing number of studies have exploited Machine Learning (ML) models to enhance and analyze MD simulations. Notably, unsupervised models that extract kinetic information from a set of parallel trajectories have been proposed, including the variational approach for Markov processes (VAMP), VAMPNets, and time-lagged variational autoencoders (TVAE). In this work, we propose a combination of adaptive sampling with active learning of kinetic models to accelerate the discovery of the conformational landscape of biomolecules. In particular, we introduce and compare several techniques that combine kinetic models with two adaptive sampling regimes (least counts and multi-agent reinforcement learning-based adaptive sampling) to enhance the exploration of conformational ensembles without introducing biasing forces. Moreover, inspired by the active learning approach of uncertainty-based sampling, we also present MaxEnt VAMPNet. This technique consists of restarting simulations from the microstates that maximize the Shannon entropy of a VAMPNet trained to perform soft discretization of metastable states. By running simulations on two test systems, the WLALL pentapeptide and the villin headpiece subdomain, we empirically demonstrate that MaxEnt VAMPNet results in faster exploration of conformational landscapes compared to the baseline and other proposed methods.

## 1 Introduction

Molecular dynamics (MD) simulations have become a widely-applied computational tool to disentangle the details of nanoscopic systems relevant to a wide range of fields, from materials engineering^1,2^ to fundamental biology.^3^ The reason for their widespread use lies in their ability to resolve the dynamics of molecular systems at excellent time and space resolutions. Nonetheless, the fine-grained time steps of MD simulations also result in high computational costs if the processes to be observed occur at long timescales. Precisely, the time interval at which an MD simulation can update atomic positions is typically restricted to the order of femtoseconds, whereas most molecular processes of interest take place at the microsecond to millisecond scale. In biology, examples of such processes include transport across transmembrane proteins,^4,5^ signal relays,^6,7^ ligand binding,^8,9^ and protein folding.^10^

A myriad of approaches have emerged to tackle the long timescale challenge in atomistic MD simulations.^11^ While numerous methods enhance the sampling of the system by modifying the potential function or the thermodynamic ensemble,^12–14^ others pursue the same goal by selectively restarting trajectories from initial conditions that favor a sampling criterion^15–17^ (see Adaptive Seeding section in ref. 11). Different problems have been studied using both types of methods, sometimes in combination, with satisfactory results.^18,19^ The choice of a suitable method will largely depend on the nature of the problem at hand. In general, adaptive seeding methods are well-suited to exploit the capabilities of large computer clusters through massively parallel simulations^20,21^ and to recover the unbiased kinetics of the system through statistical models, such as Markov state models (MSMs). ^22^

For biological systems, the thermodynamic ensembles of interest are typically isothermal-isobaric ensembles. Consequently, the probability of sampling a state decays with its energy, following a Boltzmann distribution. For this reason, one of the challenges associated with the long timescale problem in MD simulations is the sampling of rare or transition states, which are characterized by high energies and low probabilities. While biased methods can accelerate the sampling of rare states, this advantage may come at the cost of sampling an unphysical transition. In this work, we investigate the ability of unbiased simulations to explore a diverse conformational ensemble and therefore our proposed techniques fall under the adaptive seeding category.

Adaptive seeding methods can be divided into weighted ensemble and adaptive sampling techniques.^11^ Weighted ensemble techniques rely on the “splitting” and “merging” of trajectories according to their relative importance for a sampling criterion. In these methods, weights for each trajectory are tallied; splitting reduces the weight and merging increases it. These weights are introduced with the goal of recovering statistically unbiased observables. ^23^ On the other hand, adaptive sampling techniques tend to prioritize the exploration of a diverse set of molecular conformations by iteratively restarting short simulations from poorly sampled states. In past approaches, these states are obtained by using some discretization of the conformational space, such as clustering. After obtaining satisfactory coverage of the conformational landscape, the short trajectories are statistically unbiased using, for instance, MSMs.^24^ All methods presented in this study fall within the adaptive sampling category.

Machine learning (ML) is becoming critically relevant to the field of enhanced MD simulations. In particular, ML techniques have been utilized to model force fields,^25–27^ approximate optimal biasing potentials,^28,29^ and extract information from MD data.^30^ For adaptive sampling in particular, manifold learning^31,32^ and reinforcement learning (RL) algorithms^17,31,33,34^ have been applied in the past to guide simulations. Among ML methodologies, deep neural networks (DNNs) are especially promising thanks to their ability to learn arbitrarily complex nonlinear functions. ^35^

An interesting application of ML models to MD that motivates the current study is the extraction of kinetic information from a set of trajectories. The reason is that kinetics can potentially be exploited to improve the selection of initial conditions for adaptive sampling. The variational approach to Markov processes (VAMP)^36,37^ can discover an optimal mapping from input features (functions of the degrees of freedom of the system) and the slow reaction coordinates of the process. This is achieved by maximizing a variational score, often termed the VAMP score.^37^ A family of DNN models that are trained to maximize the VAMP score for a set of trajectories have also been proposed,^38,39^ with the simplest of them being termed VAMPNet.^38^ In contrast to the linear approach used in VAMP (see Feature TCCA),^37^ the DNN models can find nonlinear combinations of features and may incorporate soft state discretization as part of their architecture.^38^

*A priori*, it is unclear how to use the output of a DNN model that maximizes a VAMP score to guide adaptive sampling simulations. When fitting such a model without a discretization layer, the mapping can be interpreted as a learned embedding spanned by the slowest-changing collective variables (CVs) or reaction coordinates (RCs) of the system. This is a dimensional reduction if the output layer is smaller than the input layer. In this case, one may employ the DNN model in a similar way VAMP^36,37^ or time-lagged independent component analysis (tICA)^40–42^ are used in adaptive sampling workflows: the model projects the conformations along the slow processes and the state discretization and selection take place in this learned embedding. This is expected to improve performance when it is difficult to resolve state transitions in feature space and the number of dimensions must be reduced to achieve a reasonable clustering.

When incorporating the state discretization layer (usually realized as a softmax operation^38^) the output of these models can be interpreted as the membership probabilities of a microstate in the kinetically metastable states. Given this probabilistic interpretation, we propose incorporating an information theoretic metric to guide the choice of new restarting points for adaptive sampling. Shannon entropy (see Methods) is a metric that reveals the uncertainty of a model against the possible outcomes of an event (in this case, the uncertainty of the model in placing a microstate into a metastable state). It has been long connected to statistical mechanics^43^ and utilized to combine experimental data with MD simulations.^44–46^ In this study, we empirically show that selecting the microstates that maximize the Shannon entropy of a VAMPNet leads to improved exploration in adaptive sampling, measured as the volume of CV or tIC space observed by the generated trajectories.

We divide our study in two phases: in the initial exploratory phase, we focused on three kinetic models available in the Python library deeptime,^47^ VAMP,^37^ VAMPNets,^38^ and time-lagged variational autoencoders (TVAEs),^39^ and combined them with two adaptive sampling methods: least counts (LC) adaptive sampling and multi-agent reinforcement-learning based adaptive sampling (MA REAP).^17^ LC is a common baseline for adaptive sampling; in this technique, the states are obtained by clustering and the starting structures are selected from the clusters with the fewest members. ^48^ In MA REAP, the states are also obtained by clustering, but the starting structures are selected following a reward function that depends on the deviation of the structure with respect to the mean of the data.^17^ We compared these methods based on their ability to explore the conformational landscape of a flexible pentapeptide (sequence WLALL).^49^ Inspired by the results from these comparisons, we introduced the entropic metric for VAMPNets in the second phase of the study, where we showed that this last method achieves superior exploration when applied to two systems of different complexity: the WLALL pentapeptide and a fast-folding protein subdomain, the villin headpiece.^50^

The rest of the paper is organized as follows: the Methods section explains all the techniques introduced and compared in this paper. The Results and Discussion section is divided into three parts. In the first part, we present the results of the exploratory phase of the study. In the second part, we introduce the Shannon entropy criterion with VAMPNets for adaptive sampling and compare it with the techniques analyzed in the exploratory phase. Lastly, we compare the two most promising methods in a realistic system (the villin headpiece protein) to validate the results observed in the previous subsection. We conclude with a discussion of the advantages and limitations of our proposed methods.

## 2 Methods

### 2.1 Coupling Adaptive Sampling and Active Learning

Adaptive sampling MD is an iterative technique where the conformational landscape of a molecular system is progressively discovered by restarting simulations from poorly sampled states. Typically, the iteration is divided into two steps: running trajectories and data analysis. In the first step, trajectories initiated from the specified conformations are simulated. In the second step, the sampled points are clustered to discretize the conformational landscape into distinct states. Subsequently, a strategy is employed to determine which states to initiate the simulations anew. In the next iteration, new trajectories are executed from the selected states. This process continues until sampling is satisfactory. The selection strategy for the restarting points has a profound impact on the sampling behavior.^15,51^ Since only unbiased trajectories are typically utilized during adaptive sampling, the restarting selection strategy is the hinge point that researchers manipulate to alter the behavior of their algorithms.

Interestingly, there is a subset of ML termed active learning^52^ that realizes training as an iterative process where a model is first fit to an available data set and then new data points are queried to an oracle based on some criterion that maximally improves learning. Active learning has previously been used in combination with MD simulations to efficiently explore chemical space^53–55^ and to find model parameters.^56^ Since both adaptive sampling and active learning follow a two-stage process, we can couple both techniques. In other words, the data analysis phase of adaptive sampling becomes the learning phase of active learning. Similarly, the MD integrator acts as the oracle, so the querying phase of active learning becomes the simulation step in adaptive sampling. The information extracted by the ML model is used to select the restarting points for new simulations, which in turn will be utilized to refine the model in the next iteration. This workflow is summarized in Algorithm 1.

#### Algorithm 1

Coupled active learning–adaptive sampling

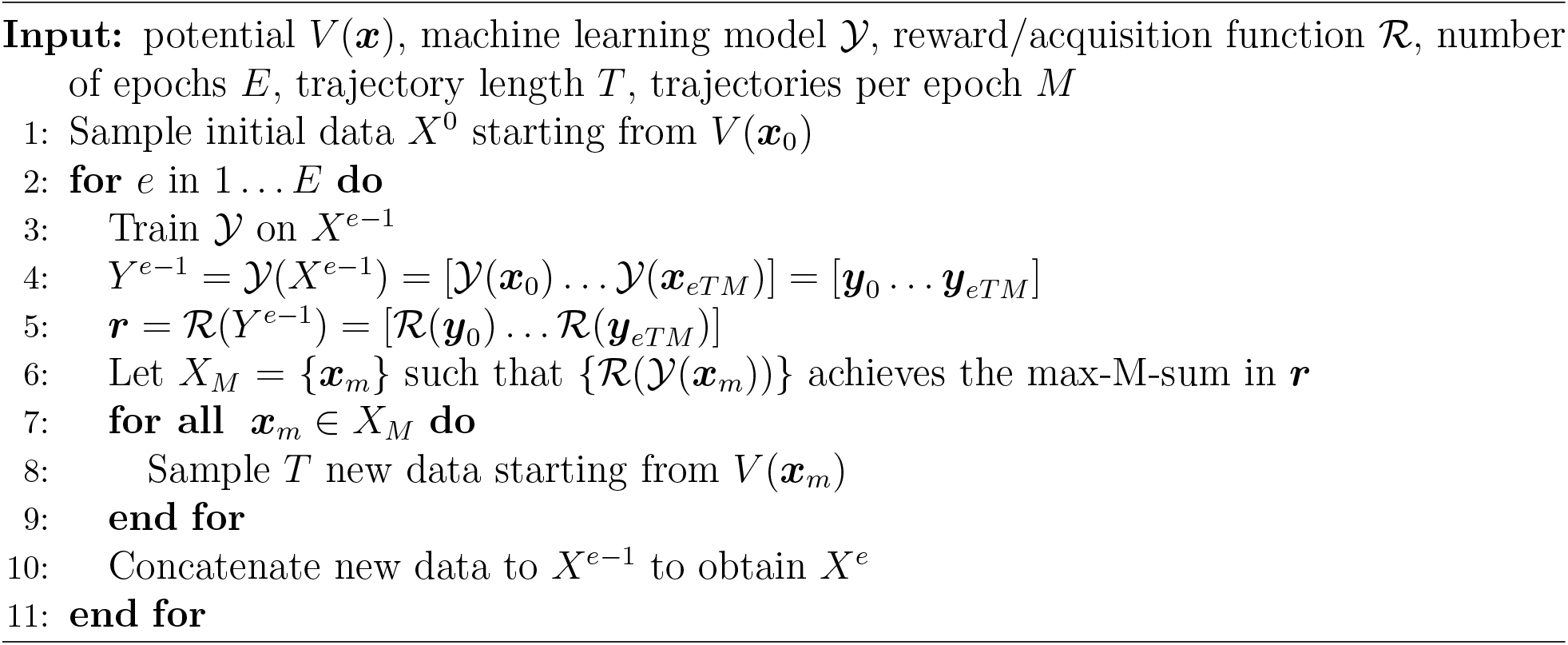

In Algorithm 1, the function ℛ outputs a scalar that is used to rank the conformations according to how desirable it is to restart simulations from them. Depending on the adaptive sampling regime employed, an additional model might be used to optimize ℛ.^17,33^ For this reason, ℛ can be interpreted as a reward function when the adaptive sampling regime depends on a RL model or an acquisition function when it does not.

### 2.2 Combining Kinetic Models with Adaptive Sampling Regimes

For Algorithm 1 to work, we must identify a suitable model *Y* that is able to extract useful information from the parallel trajectories. A relevant family of models^37–39,42^ is based on identifying the slowest processes that occur in a system. Arguably, these processes will be rate-limiting for the state transitions. The models learn the slowest processes by finding the transformations that maximize a variational score for a set of trajectories, which is usually termed the VAMP score.^37^ In mathematical terms, for a set of trajectories 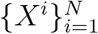 where 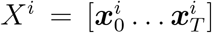, we can compute the covariance matrices *C*_00_ and *C*_11_, and time-lagged covariance matrix *C*_01_ as follows^37,38^

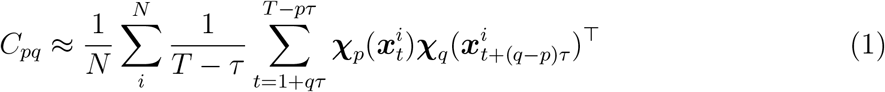

where *p* and *q* must be replaced by 0 or 1 and *τ* is the lag time measured in time steps. ***χ***_0_ and ***χ***_1_ are transformations from features or CVs to latent variables. It is possible to have ***χ***_0_ and ***χ***_1_ be identical and these transformations can be machine-learned through stochastic methods.^38^ Once these matrices have been estimated from data, they are pre-processed to remove the mean and obtain centered covariance matrices 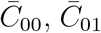 and 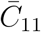. Then, a version of the VAMP score (VAMP-2)^37^ can be computed as

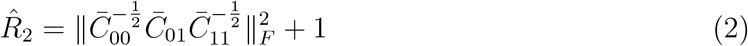

where the norm *F* is the Frobenius norm.

Different ML models have been proposed to find the transformations {*𝒳*_0_, *𝒳*_1_} and have been implemented in deeptime.^47^ Namely, the models that we employ in this study are VAMP,^37^ VAMPNets,^38^ and TVAEs.^39^ By using the VAMP-2 score as the gain function, these models learn useful mappings from the input features to the slowest-changing processes in the system.^37–39^ These learned embeddings can be used to discriminate between kinetically different states. If the output of the model has fewer dimensions than the input feature vector, then the model performs a dimensionality reduction.

In this study we use VAMP to refer to feature time-lagged canonical correlation analysis (feature TCCA).^37^ Time-lagged independent component analysis (tICA)^40,41^ and the variational approach to conformational dynamics (VAC)^36^ are subclasses of this technique.

In this case, the basis sets {𝒳_0_, 𝒳_1_} are simply the user-defined features or CVs. VAMP works by performing a truncated singular value decomposition on the matrix 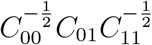to obtain *U*^*′*^*KV* ^*′*⊤^. Then, the coefficient matrices *U* and *V* are computed as 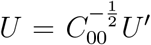 and 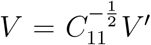. Finally, the learned projections, i.e. the left and right singular functions, can be found as 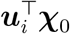 and 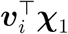 respectively, where ***u***_*i*_ and ***v***_*i*_ are the *ith* column vectors of *U* and *V*.^37,47^

In contrast to VAMP, VAMPNets are based on DNNs. They are typically implemented as multilayer perceptrons (MLPs), but other architectures have been proposed.^57^ Although in their original form VAMPNets were designed with a softmax layer for output processing to learn a soft discretization, they can also be utilized without one to learn the projection from feature space onto the slow variables of the system. There exists literature that refers to such models as “state-free” VAMPNets.^58^ The transformations {𝒳_0_, 𝒳_1_} can be stochastically learned by DNNs because the variational score in equation 2 is differentiable and thus the update gradient is well-defined.^38^

TVAEs can be conceived as extensions of VAMPNets where the DNN employed is a variational autoencoder. This is a network that consists of an encoder, which compresses the input at time *t* into a small number of dimensions or latent space. Then, the decoder reconstructs the input from the compressed dimensions at time *t* + *τ*. Since this is a time-lagged autoencoder, the encoder essentially performs a nonlinear version of TCCA. ^47^ The variational qualifier refers to the fact that the TVAE learns a probability distribution over the latent space, as opposed to simply mapping inputs to fixed points on the dimensionally-reduced coordinates. The use of variational autoencoders offers the possibility of employing the trained model for generative tasks.^47^ However, as presented in Results and Discussion, the TVAE-based sampling techniques did not perform on par with the VAMPNet-based ones.

In the exploratory phase of this study, we asked if combining such kinetic models with different adaptive sampling schemes would yield better exploration performance. We tested two adaptive sampling schemes: least counts (LC) and multi-agent reinforcement learning-based (MA REAP) adaptive sampling. While the former is a typical baseline for adaptive sampling, the latter utilizes a more complex reward function to select the restarting points for simulation.

When employing LC adaptive sampling in combination with a kinetic model, we project the conformations onto the learned embedding and then cluster the data points into discrete states. To seed the next round of simulations, we choose the centers of those clusters with the fewest number of members. In other words, ℛ is the inverse of the frame count that falls in the same cluster as the evaluated conformation.

To combine these kinetic models with MA REAP, the first steps are identical as for LC: project the conformations, cluster them, select cluster centers as the states, and select a subset of candidates based on the least counts criterion. However, when employing MA REAP, the starting structures are selected through a stakes-based reward function ^17^

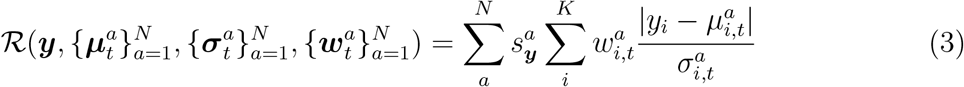

where *N* is the number of agents and *K* is the number of output dimensions. ***y*** is a cluster center or state (projected onto the embedding learned by 𝒴), and 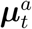 and 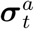 are the mean and standard deviation of the data (as estimated up to time *t*) for agent *a*. 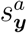 represents the stake that agent *a* has on state ***y***, which only becomes relevant when multiple agents are scouting the conformational landscape. The stake essentially determines which agent collects the data from a trajectory started at ***y*** to change its estimates of 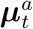 and 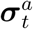.

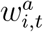 represents the weight that is assigned to a given dimension, so if a conformation displays a larger deviation along a latent variable with a higher weight, the reward is larger. The inner sum of equation 3 can be interpreted as a weighted standardized Euclidean distance. Both 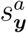 and 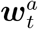 are fit from the trajectories, but 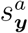 is determined at the clustering step and 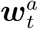 is set to maximize ℛ through quadratic optimization.^17,33^

One may ask if, in the multi-agent case, one distinct version of 𝒴 should be learned by each agent given only their own data. If this were the case, the outer sum in equation 3 would lack a meaningful interpretation since the *K* output dimensions would differ across agents. Therefore, we restrict all agents to use the same instance of a kinetic model. It might be possible to modify equation 3 to accommodate the alternative design choice, but this is out of the scope of the present study. Learning different versions of 𝒴 may alter the behavior of the algorithm by forcing agents to rely on local kinetic maps, rather than on a global model.

Utilizing the three aforementioned kinetic models with two adaptive sampling regimes yields six combinations, which are denominated {VAMP, VAMPNet, TVAE} + {LC, MA REAP} according to the model and the regime employed in each case.

### 2.3 Maximum Entropy VAMPNets

Unlike the previously described approaches, where the model *Y* projects a conformation onto the slow-changing dimensions of the system, here we are interested in VAMPNets that perform a “fuzzy clustering” by incorporating a final softmax layer. The softmax operation can be expressed as

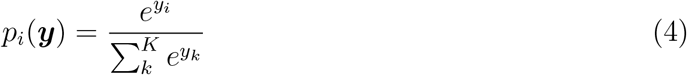

where *p*_*i*_(***y***) can be interpreted as the probability that conformation ***y*** is in the output state *i* ∈ {1, …, *K*}. For a VAMPNet, each output state corresponds to a kinetically distinct state or metastable state. Due to this probabilistic interpretation, we can compute the Shannon entropy for a given conformation,

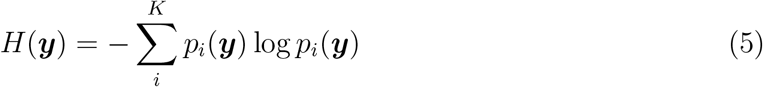

which is maximized when the predicted probability over the *K* output states is uniform or *p*_*i*_(***y***) = 1*/K* ∀*i*. In other words, *H*(***y***) is maximized when the model is uncertain about what metastable state the conformation “belongs” to. *H* can be interpreted as a choice for ℛ in Algorithm 1 when the output of 𝒴 is a probability distribution.

In the active learning community, the Shannon entropy is one of the most general and popular choices of uncertainty metric.^52^ Nonetheless, to the best of our knowledge, uncertainty sampling has never been used in combination with VAMPNets, including entropy-based sampling. In our proposed adaptive sampling technique, after 𝒴 has been fit to the data and the membership probabilities *p*_*i*_(***y***) have been obtained, the Shannon entropy, *H*(***y***), of each conformation (or a representative subset when memory becomes a concern) is computed. Then, the structures that maximize *H*(***y***) are selected to seed the next round of simulations and the workflow of Algorithm 1 proceeds. The intuition behind this method is that the VAMPNet will select structures that cannot be easily categorized, and therefore the conformations that are poorly sampled and/or lie at the interface between metastable states will be prioritized as starting simulation conditions. Figure 1 illustrates this selection criterion at play on a chaotic deterministic model, termed the Lorenz system.^59^ Here, a two-state VAMPNet was trained on a trajectory with initial conditions ***x***_0_ = (8, 7, 15)^⊤^ and default deeptime^47^ parameters for {*σ, β, ρ, h*}. When projecting the output of this VAMPNet on a different trajectory obtained with ***x***_0_ = (7, 8, 14)^⊤^, it is clearly observed that the data points that maximize the entropy (black dots) correspond to transitions between lobes (Figure 1B). In the rest of the paper we refer to this method as MaxEnt VAMPNet or MaxEnt for brevity.

**Figure 1:**
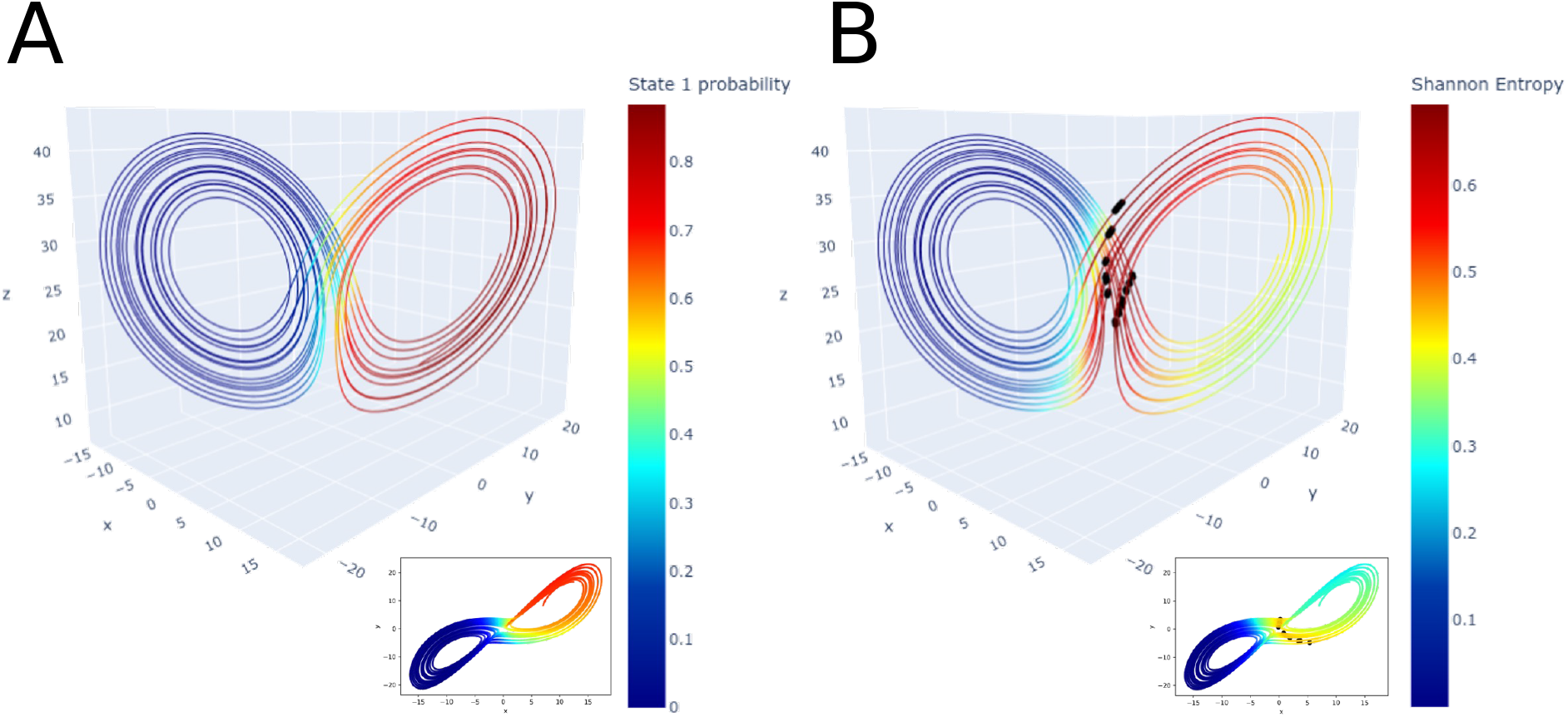
Illustration of the MaxEnt VAMPNet selection criterion in the original Lorenz system.^59^ (A) Projection of the state 1 probabilities on the validation trajectory. Inset shows projection on the *x*-*y* plane. (B) Projection of the Shannon entropy on the validation trajectory. Black dots indicate the 20 data points with the highest entropy. Inset shows projection on the *x*-*y* plane. It can be observed that the entropy maxima occur at the interface between states.

## 3 Results and Discussion

### 3.1 Uncertainty-Based Selection Criteria Achieve Superior Exploration Performance

We begin by comparing the techniques described in Section 2.2 using a pentapeptide of sequence WLALL. ^49^ This model is small enough to quickly prototype and test different methods, but it also contains a nontrivial number of degrees of freedom and slow variables. For details about the simulations, refer to the Supporting Information methods.

The input features used to fit the kinetic models were all *ϕ* and *ψ* dihedral angles (eight features in total; *ϕ*_1_ and *ψ*_5_ are undefined). The models were used to project the conformations in a two-dimensional space. For the VAMPNets, a MLP was employed with lobe duplication.^38^ The dimensions of each layer were [8, 15, 10, 10, 5, 2] with rectified linear unit (ReLu) as the activation function. These were also the dimensions of the TVAE’s encoder, while the decoder was a MLP with dimensions [2, 5, 10, 10, 15, 8]. In all cases, the lagtime was set to 20 *ps* and the batch size was 1024. We do not split the collected data into training and validation sets because our goal is to maximize the exploration rather than to validate the quality of the kinetic model. Keeping trajectories out of the analysis would preclude selecting starting structures from them (see Conclusions).

All simulations were started from the same two metastable structures obtained from a previous study.^49^ The clustering method utilized was regular space clustering (implemented in deeptime)^47^ with identical parameters in all cases (max distance = .001, max centers = 10^4^). For MA REAP, two agents with “equal” stakes and the “collaborative” regime^17^ were used. As for other MA REAP parameters, CV weights were initialized as 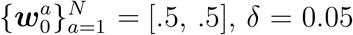, and 50 LC candidates were selected per round. ^33^ Each round consisted of 5 trajectories of 2 *ns* each and 100 training epochs of the DNN-based kinetic models. We ran all methods for 60 rounds, with 20 replicates each (total simulated time of 96 *μs*).

Figure 2 shows the results of the comparison in terms of the volume of dihedral space explored by each technique against the LC baseline. Figure 2A shows the results for the methods employing a combination of LC adaptive sampling and a kinetic model, while Figure 2B shows the same for the techniques involving MA REAP. Figure 2A shows that combining LC adaptive sampling with a VAMPNet produced a considerable advantage against the LC baseline (100% increase in explored volume after 600 *ns*). On the other hand, TVAE + LC only yielded an advantage of approximately 25%, but for *t <* 600 *ns* the difference is smaller and not statistically significant. VAMP + LC did not yield a statistically significant advantage for the length of simulations tested. The results from Figure 2B show that MA REAP increased the explored volume by approximately 40% after 60 rounds, but in this case the use of the kinetic models only produced marginal gains. The difference in performance between MA REAP and the combination of {VAMP, VAMPNet, TVAE} + MA REAP was not statistically significant.

**Figure 2:**
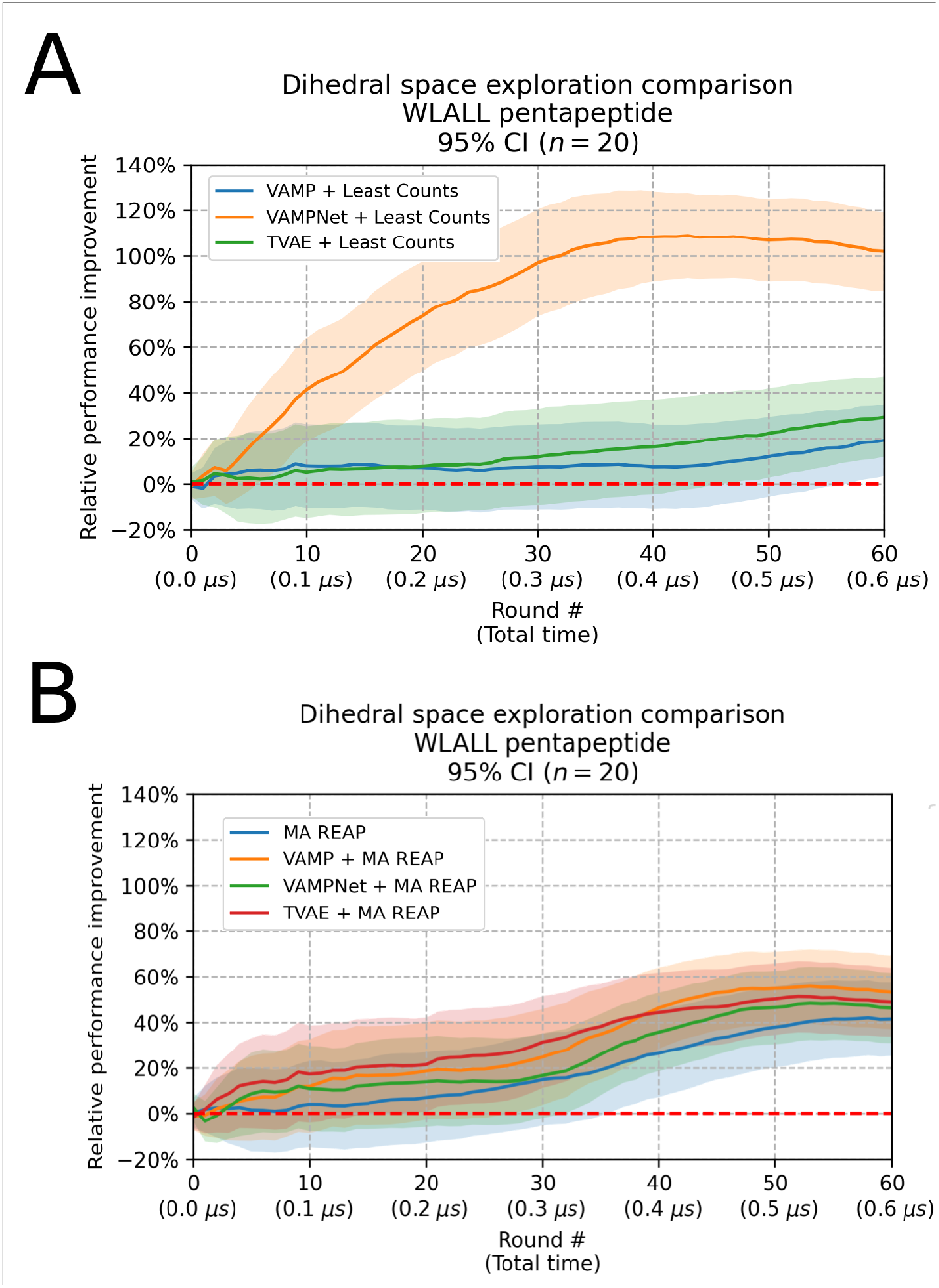
Relative increase in dihedral space volume explored on WLALL pentapeptide across techniques. The same baseline, LC adaptive sampling with no kinetic model, was used in both plots (dashed line). Curves show mean for 20 replicates with a 95% CI. (A) Comparison with {VAMP, VAMPNet, TVAE} + LC. (B) Comparison with MA REAP and {VAMP, VAMPNet, TVAE} + MA REAP.

Figure 3 provides a visual depiction of dihedral space exploration by the baseline (LC) and the best method (VAMPNet + LC). The figure shows the Ramachandran plots for the three central amino acids in the WLALL peptide for the first three replicates (for all replicates see Figures S1–S4). These plots show that the states with *ϕ*_2_ *>* 0 were not thoroughly explored by LC, whereas most replicates for VAMPNet + LC discovered this portion of the landscape. The first point of interest raised by these results is the success of VAMPNet + LC at accelerating exploration, even when compared against VAMPNet + MA REAP, which uses the same kinetic model and a more sophisticated adaptive sampling regime. This result can be explained by the fact that the reward function in MA REAP (equation 3) relies on a distance metric between the states and the distribution mean. Since the kinetic model is fit with a data set that incorporates new trajectories after each round, the mapping from feature space to latent space changes, distorting distances and, consequently, the deviations of the states. This is likely to result in an inefficient estimation of the weights that MA REAP utilizes to prioritize a given direction in exploration. Thus, we observe poor performance gains from combining MA REAP and a kinetic model. Another important comparison to make is that between VAMPNet + LC and {VAMP, TVAE} + LC. In this case, the same adaptive sampling regime is used, but the kinetic model changes. The poor performance of VAMP + LC against VAMPNet + LC arises from the fact that a linear method is inefficient at discriminating between kinetically distinct states because the boundaries are not linear in dihedral space. On the other hand, TVAE is also based on a DNN and the encoder can learn a latent space that separates the metastable states. However, training a variational autoencoder is more demanding than training a MLP, as the TVAE must learn the probability distribution over the latent space and a decoder must be simultaneously fit. Although it remains interesting to utilize models such as TVAEs (which allow for generative inference) in future applications, in this study we limit ourselves to observe that they do not accelerate adaptive sampling at a similar rate as simpler MLP-based models (VAMPNets).

**Figure 3:**
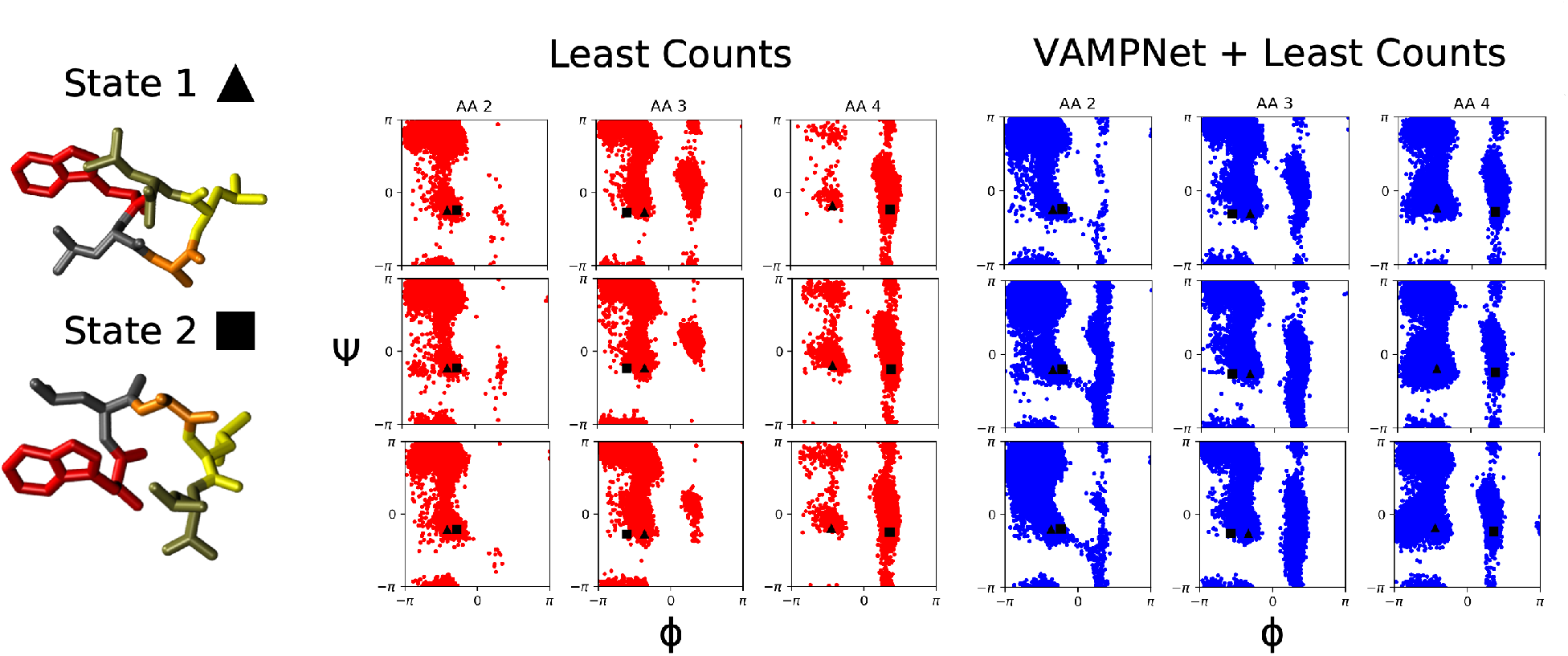
Ramachandran plots for central amino acids in WLALL peptide for baseline method (LC) and most successful method (VAMPNet + LC). Each row corresponds to a different replicate, each column to a different amino acid. (Left) Initial states employed in all simulations. These conformations were projected on the plots. (Center) Ramachandran plots for LC. (Right) Ramachandran plots for VAMPNet + LC.

Overall, in this section we showed that VAMPNet + LC achieved an advantage of ∼100% against the LC baseline and ∼60% against MA REAP methods in the pentapeptide model. This observation motivated the steps taken in the following section.

### 3.2 MaxEnt Achieves Faster Discovery of the Conformational Landscape

Inspired by the results from the previous section, we further investigated the origin of the large advantage displayed by VAMPNet + LC. In general, LC is better than continuous MD simulations because it prioritizes the sampling of poorly characterized states. We hypothesized that a determining factor in the success of VAMPNet + LC is the selection of data points that lead to “informative” trajectories for the VAMPNet. In other words, LC selected structures that resulted in better training examples for the DNN model, allowing a productive separation of states in the latent space, and thus encouraged the discovery of new regions of the conformational landscape in future iterations. If this is the case, then another selection regime that queries data points to maximize learning should result in advantageous exploration, even if that regime is not related to a known adaptive sampling technique. For this reason, we decided to utilize entropy-based sampling, which is a common choice in the active learning community (see Methods), and termed the new method MaxEnt VAMPNet. In this method, the output of the VAMPNet is not interpreted as coordinates in latent space, but rather as membership probabilities in the output states. For this reason, we used a larger output layer (eight states) and included a softmax layer as the final operation. The number of parameters in the hidden layers were also increased, the new dimensions were [8, 16, 32, 64, 128, 256, 128, 64, 32, 16, 8]. New runs with VAMPNet + LC were performed with identical layer sizes to observe the effect of increasing the number of parameters. The number of training epochs per round was kept at 100, but the batch size was increased to 2048. Other details were identical as the previous section. The length of trajectories, number of rounds, and number of replicates were also kept identical for a total simulated time of 36 *μs*.

Figure 4 shows the results from this trial. The VAMPNet + LC result from the previous section is also plotted in Figure 4A for clarity. This plot shows that MaxEnt reached the same level of performance improvement as VAMPNet + LC (with the smaller DNN from the previous section). However, the entropy-based method reached this level of advantage after only ∼150 *ns* instead of the ∼300 *ns* that it took VAMPNet + LC. It is important to observe that using a larger VAMPNet with more output dimensions harms the performance of VAMPNet + LC. This is likely due to the fact that the quality of clustering degrades when using eight output dimensions instead of two. This highlights an advantage of MaxEnt, as this technique does not rely on clustering and state assignment is handled directly by the VAMPNet. For a projection of the data from all replicates of MaxEnt on Ramachandran plots, see Figures S5–S6.

**Figure 4:**
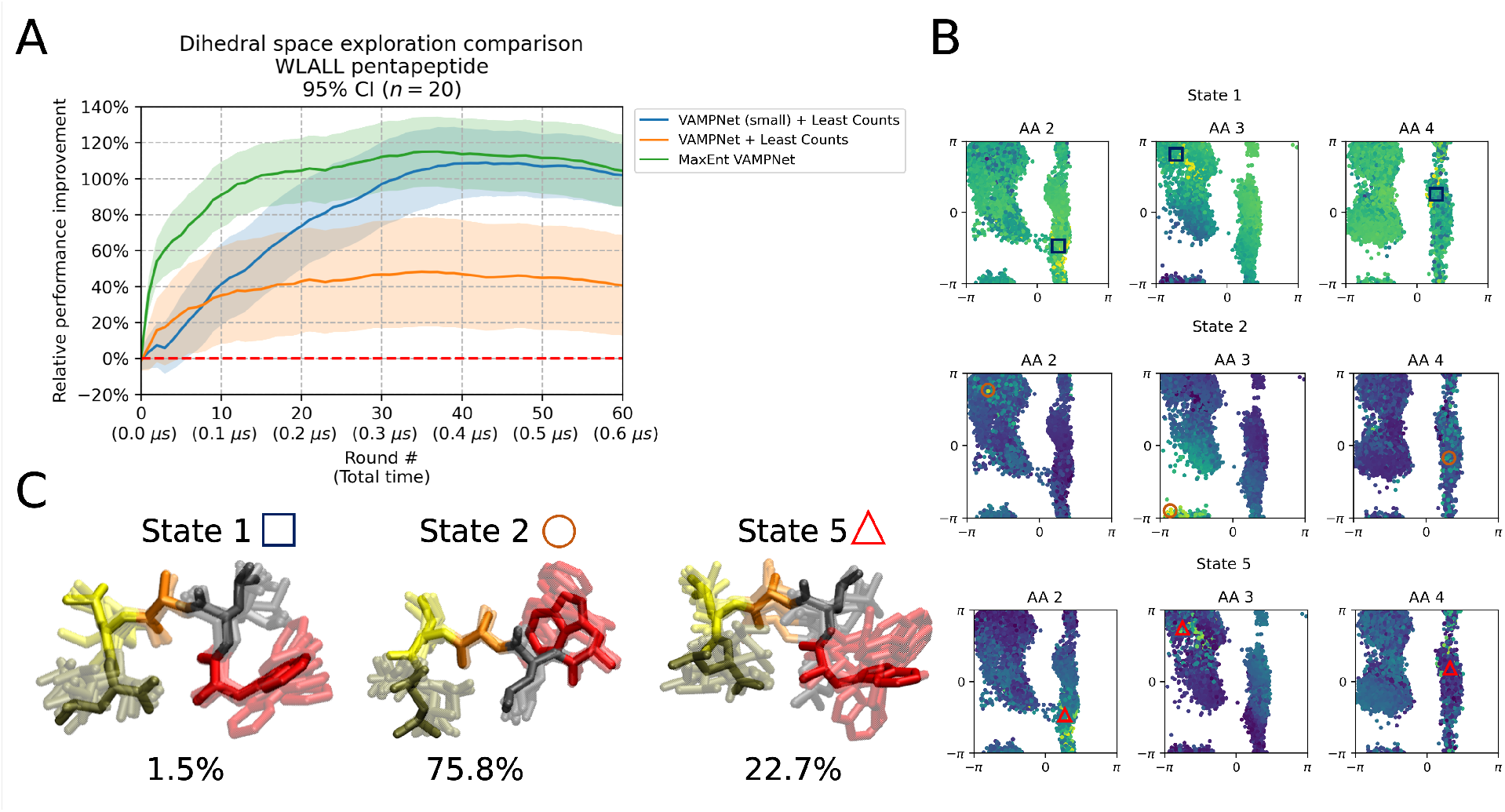
Results for comparison involving MaxEnt VAMPNet. (A) Relative increase in dihedral space volume explored on WLALL peptide across techniques. LC adaptive sampling with no kinetic model (dashed line), was used as baseline. Curves show mean for 20 replicates with a 95% CI. (B) Projection of the three VAMPNet states with nonempty populations on the Ramachandran plots of L2-A3-L4 from the first replicate of MaxEnt. (C) Conformations that maximize the probability for the output states 1, 2, and 5 from (B). Percentage populations are shown below each state.

Although we do not expect MaxEnt to produce a fully validated kinetic model (see Conclusions), we inspected the output of the VAMPNet obtained by this technique. Figure 4B shows the projection on Ramachandran plots of the only three states with nonempty memberships obtained from the first replicate of MaxEnt. In general, we observe gradients that indicate that the VAMPNet has learned a useful separation of states in dihedral space. However, state 5 shows very diverse conformations, unlike states 1 and 2 (Figure 4C), showing that the model could be lumping unrelated conformations into the same state.

In summary, in this section we showed that MaxEnt can achieve the same advantage as VAMPNet + LC in a shorter amount of time, suggesting that the former option is a more favorable choice of adaptive sampling regime. In the following section, we compare these two techniques in a more realistic model to assess if the trends observed in the pentapeptide model translate to a larger protein.

### 3.3 MaxEnt Shows an Advantage in a Realistic System

The villin headpiece subdomain (PDB ID: 1YRF)^60^ is a 35-amino acid, fast-folding protein that represents a more realistic system for MD simulations. The input features used were all pairwise *C*_*α*_ distances (separated by at least two residues). Therefore, we obtain 528 features. Since the number of features is too large to produce a reasonable clustering for LC without applying dimensionality reduction techniques, we drop this baseline and instead use VAMPNet + LC as the standard to assess the performance of MaxEnt. According to the results from the previous sections, this new baseline is significantly more demanding than vanilla LC. In all cases, the dimensions of the VAMPNets were [528, 512, 256, 128, 64, 32, 16, 8]. Batch size was set to 1024 and lobe duplication was employed. Lagtime used was 100 *ps*. Each round consisted of 10 trajectories of 10 *ns* each and 100 training epochs for the VAMPNet. We performed 10 replicates per method with 10 rounds per replicate (total simulated time of 20 *μs*). Other details were identical to previous sections. For details about the MD simulations, refer to the Supplementary Information methods.

Since it is impractical to compute the explored volume in a 528-dimensional space, we pool all the data from both methods and fit a VAMP model to project the trajectories onto a common 8-dimensional tIC space. We then compute the explored volume in this space. Figure 5 shows the comparison for VAMPNet + LC vs. MaxEnt. We can observe that there are no statistically significant differences between both methods until *t* = 700 *ns*. After 1 *μs*, MaxEnt shows an average exploration advantage of ∼50% with a 95% confidence interval of approximately [20%, 90%]. The volume explored by individual replicates is plotted in Figure S7 to easily observe the distribution for each method.

**Figure 5:**
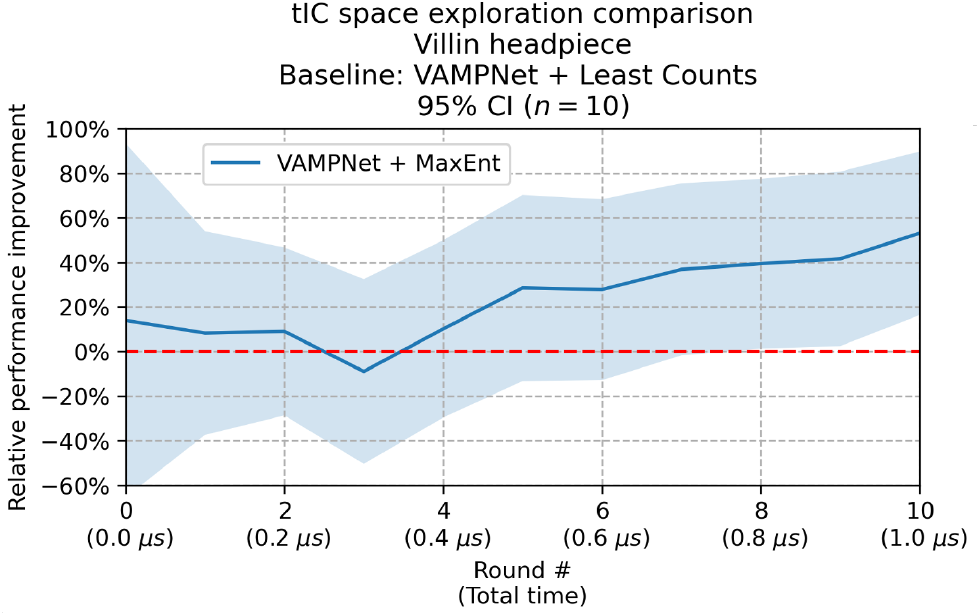
Relative increase in tIC space volume explored for MaxEnt simulations of the villin headpiece subdomain. VAMPNet + LC (dashed line), was used as baseline. The curve shows the mean for 10 replicates with a 95% CI.

Figure 6A shows the tIC1-tIC2 landscapes for the first replicate of each method and for a single continuous trajectory of the same total length. The landscapes for all replicates are available in Figures S8–S11. While 1 *μs* simulations are insufficient to observe unfolding at *T* = 300 *K*, we can observe differences in the conformational ensemble explored by MaxEnt since it discovers an area of the landscape that remains uncharted by the continuous trajectory and VAMPNet + LC.

**Figure 6:**
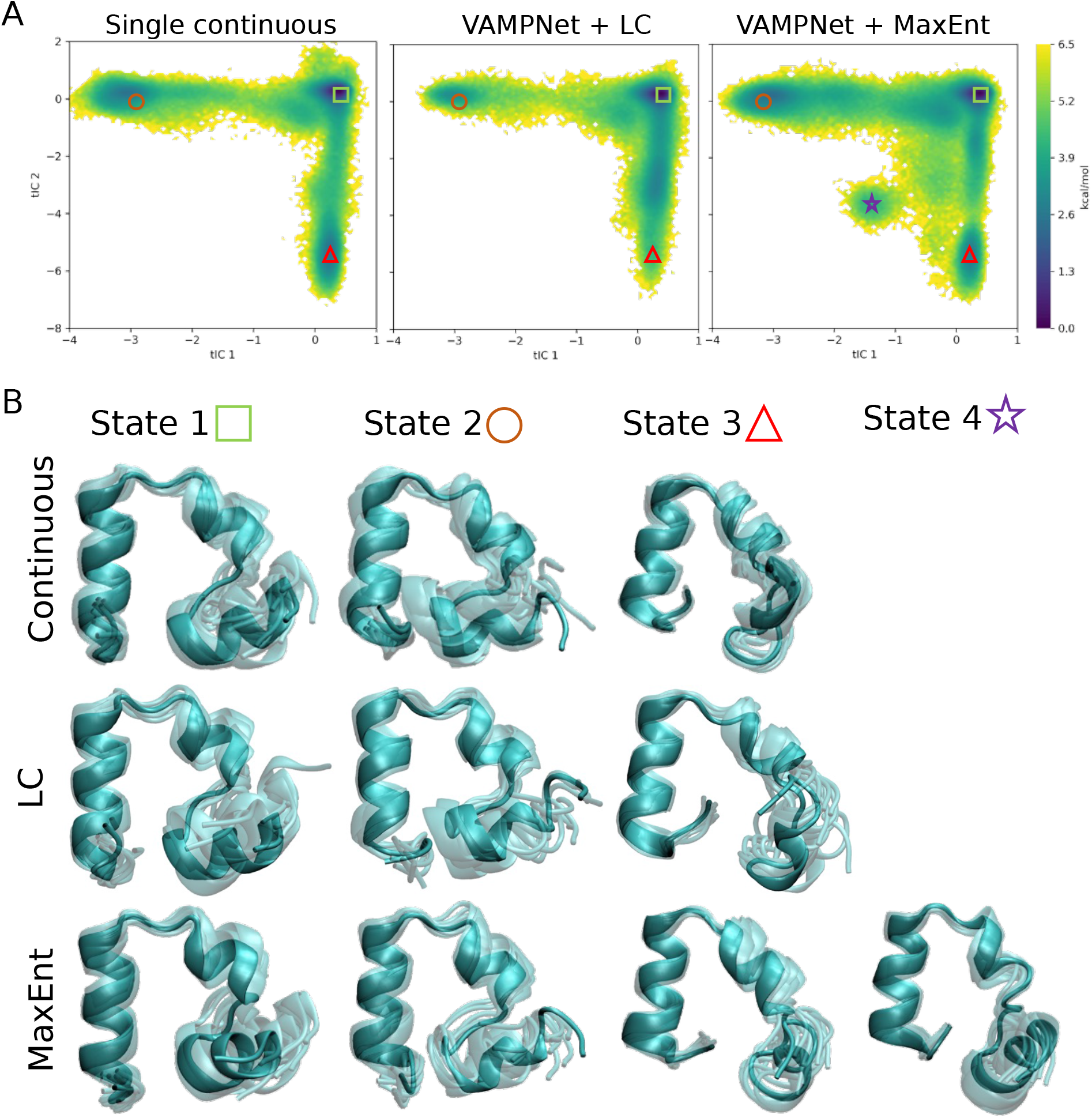
(A) tIC1-tIC2 landscapes for villin headpiece corresponding to the first replicate of each method. (B) Representative conformations of different states discovered by each technique.

Figure 6B shows a representative set of conformations discovered by each technique. State 1 represents the native folded structure, state 2 is a “closed” conformation where the C-terminus interacts with F51 and G52, state 3 is a conformation where the N-terminal *α*-helix bends perpendicularly to the plane spanned by the two other *α*-helices, and state 4 (often observed by MaxEnt but not by VAMPNet + LC) is an intermediate state between 2 and 3. The difference in explored area can be explained by the ability of MaxEnt to discover state 4 with a higher chance than VAMPNet + LC.

In summary, MaxEnt showed a statistically significant advantage in exploration against the challenging VAMPNet + LC baseline in a realistic system, indicating that entropy-based sampling is likely to be a better choice between these two techniques. Due to the long timescale of villin unfolding,^10^ neither method reached a denatured conformation, but this was expected for the simulation conditions.

## 4 Conclusions

In this study, we proposed new techniques involving active learning of DNN-based kinetic models to accelerate exploration in adaptive sampling MD simulations. Our results show that entropy-based sampling of a VAMPNet achieves fastest exploration of the conformational landscape in both simulated systems. Besides showing a better exploration behavior, the VAMPNet grants MaxEnt the convenience of skipping the clustering step altogether. This eliminates design decisions because clustering parameters must be set. Using a suboptimal set of clustering parameters in adaptive sampling can frustrate the rate of exploration or obfuscate subsequent data analysis.

However, MaxEnt also suffers from limitations. Training a VAMPNet in each iteration of adaptive sampling can be time consuming and computationally intensive. Nonetheless, this task is expected to represent a small fraction of the computational expense in adaptive sampling because MD simulations of large systems remain slow in comparison. Another challenge in the use of VAMPNets for adaptive sampling is their validation. Arguably, a researcher has a few options to validate the model at each sampling iteration: (1) set aside some trajectories to use as an uncorrelated validation set, (2) exclude some random {***x***_*t*_, ***x***_*t*+*τ*_ } pairs to use as a (correlated) validation set, or (3) use some *k*-fold cross validation approach. All options have advantages and downsides. For (1), setting aside entire trajectories can harm the exploration rate since the validation conformations cannot be selected to restart simulations. However, this method ensures that the validation set is uncorrelated with the training set, so the validation score is more reliable. In (2), the opposite is the case. Since the validation structures are correlated to those used in training, the user can select conformations that are similar to those withheld, but the validation score may not be indicative of the true performance of the model. Finally, (3) can produce an unbiased validation score if the data is divided into uncorrelated groups. After applying *k*-fold cross validation, the model can be retrained on the entire data set to select new structures for simulation. However, this alternative is computationally expensive in comparison to (1) or (2). Validating the model may alert the researcher that the VAMPNet employed is underfitting or overfitting the data and, thus, the model must be modified before proceeding. This represents an onerous effort that is not found in other adaptive sampling algorithms. In our trials, we decided to test our techniques without submitting the kinetic models to validation and observe if an exploration advantage was achieved regardless. Future work will involve studying the relationship between validation scores and exploration performance. It must be noted that entropy-based sampling is expected to provide robustness to MaxEnt because this uncertainty metric inherently makes the least assumptions about the model’s knowledge.^52^

Although it is regular practice among adaptive sampling MD practitioners to use tICA^40,41^ to reduce the dimensionality of the input features before applying a selection criterion to restart simulations, our tests involving similar approaches (i.e., VAMP + {LC, MA REAP}) did not yield promising results in comparison to VAMPNet + {LC, MaxEnt}. Nonetheless, this does not preclude that, for other systems, an advantage might be achieved by employing VAMP or tICA in combination with LC instead of applying LC on the feature space. In terms of applying MaxEnt in practice, the common workflow of adaptive sampling is not radically altered. In all adaptive sampling schemes, the data must be centralized at some point to perform the analysis step. This step can be replaced by the fitting of a VAMPNet (or an ensemble of VAMPNets for the sake of robustness) on the collected data. Once that the starting conformations are selected, the trajectories can be run in decentralized clusters as usual.

Lastly, it is important to note that, to the best of our knowledge, there is no theoretical foundation that indicates that the VAMP-2 gain function is the optimal choice to accelerate adaptive sampling through entropy-based sampling. This choice is intuitive because a VAMPNet trained to maximize this score learns to discriminate between metastable states,^38^ and therefore the conformations that maximize the Shannon entropy are more likely to be low-probability and/or poorly-sampled structures. Moreover, there are several other active learning approaches besides entropy-based sampling^52^ that have not been explored in this work. Future studies in this direction will explore questions of the optimal choice of loss function and active learning regime for the purposes of adaptive sampling.

## Acknowledgement

The authors acknowledge support from the National Science Foundation Early CAREER Award (NSF MCB-1845606).

## Supporting Information Available

Supporting methods and figures. Code necessary to reproduce the simulations is available on https://github.com/ShuklaGroup/MaxEntVAMPNet.

## Notes

### Competing Interest Statement

The authors have declared no competing interest.

## References

(1) Lau, D.; Jian, W.; Yu, Z.; Hui, D. Nano-engineering of construction materials using molecular dynamics simulations: Prospects and challenges. Composites Part B: Engineering 2018, 143, 282–291.

(2) Jackson, N. E. Coarse-graining organic semiconductors: the path to multiscale design. The Journal of Physical Chemistry B 2020, 125, 485–496.

(3) Weigle, A. T.; Feng, J.; Shukla, D. Thirty years of molecular dynamics simulations on posttranslational modifications of proteins. Physical Chemistry Chemical Physics 2022,

(4) Chan, M. C.; Selvam, B.; Young, H. J.; Procko, E.; Shukla, D. The substrate import mechanism of the human serotonin transporter. Biophysical journal 2022, 121, 715– 730.

(5) Feng, J.; Selvam, B.; Shukla, D. How do antiporters exchange substrates across the cell membrane? An atomic-level description of the complete exchange cycle in NarK. Structure 2021, 29, 922–933.

(6) Shukla, D.; Meng, Y.; Roux, B.; Pande, V. S. Activation pathway of Src kinase reveals intermediate states as targets for drug design. Nature communications 2014, 5, 1–11.

(7) Kohlhoff, K. J.; Shukla, D.; Lawrenz, M.; Bowman, G. R.; Konerding, D. E.; Belov, D.; Altman, R. B.; Pande, V. S. Cloud-based simulations on Google Exacycle reveal ligand modulation of GPCR activation pathways. Nature chemistry 2014, 6, 15–21.

(8) Shan, Y.; Kim, E. T.; Eastwood, M. P.; Dror, R. O.; Seeliger, M. A.; Shaw, D. E. How does a drug molecule find its target binding site? Journal of the American Chemical Society 2011, 133, 9181–9183.

(9) Chen, J.; White, A.; Nelson, D. C.; Shukla, D. Role of substrate recognition in modulating strigolactone receptor selectivity in witchweed. Journal of Biological Chemistry 2021, 297.

(10) Lindorff-Larsen, K.; Piana, S.; Dror, R. O.; Shaw, D. E. How fast-folding proteins fold. Science 2011, 334, 517–520.

(11) Hénin, J.; Lelièvre, T.; Shirts, M.; Valsson, O.; Delemotte, L. Enhanced Sampling Methods for Molecular Dynamics Simulations [Article v1. 0]. Living Journal of Computational Molecular Science 2022, 4, 1583–1583.

(12) Hamelberg, D.; Mongan, J.; McCammon, J. A. Accelerated molecular dynamics: a promising and efficient simulation method for biomolecules. The Journal of chemical physics 2004, 120, 11919–11929.

(13) Yu, T.-Q.; Lu, J.; Abrams, C. F.; Vanden-Eijnden, E. Multiscale implementation of infinite-swap replica exchange molecular dynamics. Proceedings of the National Academy of Sciences 2016, 113, 11744–11749.

(14) Laio, A.; Rodriguez-Fortea, A.; Gervasio, F. L.; Ceccarelli, M.; Parrinello, M. Assessing the accuracy of metadynamics. The journal of physical chemistry B 2005, 109, 6714– 6721.

(15) Zimmerman, M. I.; Porter, J. R.; Sun, X.; Silva, R. R.; Bowman, G. R. Choice of adaptive sampling strategy impacts state discovery, transition probabilities, and the apparent mechanism of conformational changes. Journal of chemical theory and computation 2018, 14, 5459–5475.

(16) Zuckerman, D. M.; Chong, L. T. Weighted ensemble simulation: review of methodology, applications, and software. Annual review of biophysics 2017, 46, 43.

(17) Kleiman, D. E.; Shukla, D. Multiagent reinforcement learning-based adaptive sampling for conformational dynamics of proteins. Journal of Chemical Theory and Computation 2022, 18, 5422–5434.

(18) Moffett, A. S.; Bender, K. W.; Huber, S. C.; Shukla, D. Molecular dynamics simulations reveal the conformational dynamics of Arabidopsis thaliana BRI1 and BAK1 receptor-like kinases. Journal of Biological Chemistry 2017, 292, 12643–12652.

(19) Zhao, C.; Shukla, D. Molecular basis of the activation and dissociation of dimeric PYL2 receptor in abscisic acid signaling. Physical Chemistry Chemical Physics 2022, 24, 724– 734.

(20) Zimmerman, M. I.; Porter, J. R.; Ward, M. D.; Singh, S.; Vithani, N.; Meller, A.; Mallimadugula, U. L.; Kuhn, C. E.; Borowsky, J. H.; Wiewiora, R. P., et al. SARS-CoV-2 simulations go exascale to predict dramatic spike opening and cryptic pockets across the proteome. Nature chemistry 2021, 13, 651–659.

(21) Russo, J. D.; Zhang, S.; Leung, J. M.; Bogetti, A. T.; Thompson, J. P.; DeGrave, A. J.; Torrillo, P. A.; Pratt, A.; Wong, K. F.; Xia, J., et al. WESTPA 2.0: High-performance upgrades for weighted ensemble simulations and analysis of longer-timescale applications. Journal of Chemical Theory and Computation 2022, 18, 638–649.

(22) Chan, M. C.; Shukla, D. Markov state modeling of membrane transport proteins. Journal of Structural Biology 2021, 213, 107800.

(23) Aristoff, D.; Copperman, J.; Simpson, G.; Webber, R.; Zuckerman, D. Weighted ensemble: Recent mathematical developments. arXiv preprint arXiv:2206.14943 2022,

(24) Husic, B. E.; Pande, V. S. Markov state models: From an art to a science. Journal of the American Chemical Society 2018, 140, 2386–2396.

(25) Blank, T. B.; Brown, S. D.; Calhoun, A. W.; Doren, D. J. Neural network models of potential energy surfaces. The Journal of chemical physics 1995, 103, 4129–4137.

(26) Smith, J. S.; Nebgen, B. T.; Zubatyuk, R.; Lubbers, N.; Devereux, C.; Barros, K.; Tretiak, S.; Isayev, O.; Roitberg, A. E. Approaching coupled cluster accuracy with a general-purpose neural network potential through transfer learning. Nature communications 2019, 10, 1–8.

(27) Gkeka, P.; Stoltz, G.; Barati Farimani, A.; Belkacemi, Z.; Ceriotti, M.; Chodera, J. D.; Dinner, A. R.; Ferguson, A. L.; Maillet, J.-B.; Minoux, H., et al. Machine learning force fields and coarse-grained variables in molecular dynamics: application to materials and biological systems. Journal of chemical theory and computation 2020, 16, 4757–4775.

(28) Wang, D.; Wang, Y.; Chang, J.; Zhang, L.; Wang, H.; E., W. Efficient sampling of high-dimensional free energy landscapes using adaptive reinforced dynamics. Nature Computational Science 2021, 2, 20–29.

(29) Guo, A. Z.; Sevgen, E.; Sidky, H.; Whitmer, J. K.; Hubbell, J. A.; de Pablo, J. J. Adaptive enhanced sampling by force-biasing using neural networks. The Journal of Chemical Physics 2018, 148, 134108.

(30) Glielmo, A.; Husic, B. E.; Rodriguez, A.; Clementi, C.; Nóe, F.; Laio, A. Unsupervised Learning Methods for Molecular Simulation Data. Chemical Reviews 2021, 121, 9722– 9758.

(31) Buenfil, J.; Koelle, S. J.; Meila, M. Tangent Space Least Adaptive Clustering. ICML 2021 Workshop on Unsupervised Reinforcement Learning. 2021.

(32) Preto, J.; Clementi, C. Fast recovery of free energy landscapes via diffusion-map-directed molecular dynamics. Phys. Chem. Chem. Phys. 2014, 16, 19181–19191.

(33) Shamsi, Z.; Cheng, K. J.; Shukla, D. Reinforcement Learning Based Adaptive Sampling: REAPing Rewards by Exploring Protein Conformational Landscapes. The Journal of Physical Chemistry B 2018, 122, 8386–8395.

(34) Pérez, A.; Herrera-Nieto, P.; Doerr, S.; Fabritiis, G. D. AdaptiveBandit: A Multiarmed Bandit Framework for Adaptive Sampling in Molecular Simulations. Journal of Chemical Theory and Computation 2020, 16, 4685–4693.

(35) Hornik, K.; Stinchcombe, M.; White, H. Multilayer feedforward networks are universal approximators. Neural Networks 1989, 2, 359–366.

(36) Nüske, F.; Keller, B. G.; Pérez-Hernández, G.; Mey, A. S. J. S.; Noé, F. Variational Approach to Molecular Kinetics. Journal of Chemical Theory and Computation 2014, 10, 1739–1752.

(37) Wu, H.; Noé, F. Variational Approach for Learning Markov Processes from Time Series Data. Journal of Nonlinear Science 2019, 30, 23–66.

(38) Mardt, A.; Pasquali, L.; Wu, H.; Noé, F. VAMPnets for deep learning of molecular kinetics. Nature Communications 2018, 9.

(39) Wehmeyer, C.; Noé, F. Time-lagged autoencoders: Deep learning of slow collective variables for molecular kinetics. The Journal of Chemical Physics 2018, 148, 241703.

(40) Schwantes, C. R.; Pande, V. S. Improvements in Markov state model construction reveal many non-native interactions in the folding of NTL9. Journal of chemical theory and computation 2013, 9, 2000–2009.

(41) Pérez-Hernández, G.; Paul, F.; Giorgino, T.; De Fabritiis, G.; Noé, F. Identification of slow molecular order parameters for Markov model construction. The Journal of chemical physics 2013, 139, 07B604 1.

(42) Wu, H.; Nüske, F.; Paul, F.; Klus, S.; Koltai, P.; Noé, F. Variational Koopman models: Slow collective variables and molecular kinetics from short off-equilibrium simulations. The Journal of Chemical Physics 2017, 146, 154104.

(43) Jaynes, E. T. Information Theory and Statistical Mechanics. Physical Review 1957, 106, 620–630.

(44) Bottaro, S.; Bengtsen, T.; Lindorff-Larsen, K. Methods in Molecular Biology; Springer US, 2020; pp 219–240.

(45) Boomsma, W.; Ferkinghoff-Borg, J.; Lindorff-Larsen, K. Combining Experiments and Simulations Using the Maximum Entropy Principle. PLoS Computational Biology 2014, 10, e1003406.

(46) Amirkulova, D. B.; White, A. D. Recent advances in maximum entropy biasing techniques for molecular dynamics. Molecular Simulation 2019, 45, 1285–1294.

(47) Hoffmann, M.; Scherer, M.; Hempel, T.; Mardt, A.; de Silva, B.; Husic, B. E.; Klus, S.; Wu, H.; Kutz, N.; Brunton, S. L., et al. Deeptime: a Python library for machine learning dynamical models from time series data. Machine Learning: Science and Technology 2021, 3, 015009.

(48) Bowman, G. R.; Ensign, D. L.; Pande, V. S. Enhanced Modeling via Network Theory: Adaptive Sampling of Markov State Models. Journal of Chemical Theory and Computation 2010, 6, 787–794.

(49) Scherer, M. K.; Trendelkamp-Schroer, B.; Paul, F.; Pérez-Hernández, G.; Hoffmann, M.; Plattner, N.; Wehmeyer, C.; Prinz, J.-H.; Noé, F. PyEMMA 2: A Software Package for Estimation, Validation, and Analysis of Markov Models. Journal of Chemical Theory and Computation 2015, 11, 5525–5542.

(50) Žoldak, G.; Stigler, J.; Pelz, B.; Li, H.; Rief, M. Ultrafast folding kinetics and cooperativity of villin headpiece in single-molecule force spectroscopy. Proceedings of the National Academy of Sciences 2013, 110, 18156–18161.

(51) Hruska, E.; Abella, J. R.; Nüske, F.; Kavraki, L. E.; Clementi, C. Quantitative comparison of adaptive sampling methods for protein dynamics. The Journal of Chemical Physics 2018, 149, 244119.

(52) Settles, B. Active Learning ; Springer International Publishing, 2012.

(53) Smith, J. S.; Nebgen, B.; Lubbers, N.; Isayev, O.; Roitberg, A. E. Less is more: Sampling chemical space with active learning. The Journal of chemical physics 2018, 148, 241733.

(54) Shmilovich, K.; Mansbach, R. A.; Sidky, H.; Dunne, O. E.; Panda, S. S.; Tovar, J. D.; Ferguson, A. L. Discovery of self-assembling π-conjugated peptides by active learning-directed coarse-grained molecular simulation. The Journal of Physical Chemistry B 2020, 124, 3873–3891.

(55) Thompson, J.; Walters, W. P.; Feng, J. A.; Pabon, N. A.; Xu, H.; Goldman, B. B.; Moustakas, D.; Schmidt, M.; York, F. Optimizing Active Learning for Free Energy Calculations. Artificial Intelligence in the Life Sciences 2022, 100050.

(56) Lindsey, R. K.; Fried, L. E.; Goldman, N.; Bastea, S. Active learning for robust, high-complexity reactive atomistic simulations. The Journal of Chemical Physics 2020, 153, 134117.

(57) Ghorbani, M.; Prasad, S.; Klauda, J. B.; Brooks, B. R. GraphVAMPNet, using graph neural networks and variational approach to Markov processes for dynamical modeling of biomolecules. The Journal of Chemical Physics 2022, 156, 184103.

(58) Chen, W.; Sidky, H.; Ferguson, A. L. Nonlinear discovery of slow molecular modes using state-free reversible VAMPnets. The Journal of chemical physics 2019, 150, 214114.

(59) Lorenz, E. N. Deterministic nonperiodic flow. Journal of atmospheric sciences 1963, 20, 130–141.

(60) Chiu, T. K.; Kubelka, J.; Herbst-Irmer, R.; Eaton, W. A.; Hofrichter, J.; Davies, D. R. High-resolution x-ray crystal structures of the villin headpiece subdomain, an ultrafast folding protein. Proceedings of the National Academy of Sciences 2005, 102, 7517–7522.

