## Supporting Information for "Active Learning of the Conformational Ensemble of Proteins using Maximum Entropy VAMPNets"

### 1 Supporting Methods

#### 1.1 All-atom Molecular Dynamics Simulations

All-atom MD simulations were carried out in OpenMM 7.7.[S1] The Langevin integrator was configured to use a temperature of 300 K, a friction coefficient of 1 $ps^{-1}$, and a time step of 2 fs. The force field used in all cases was Amber ff14SB.[S2] For computational speed, the generalized Born model (GBn2) for implicit solvent was utilized. Consequently, no barostat or periodic boundary conditions were set. All bonds involving hydrogen were constrained.

For the WLALL pentapeptide, the initial PDB structures were obtained from ref. S3. The structure was processed with tleap from AmberTools22. Frames were saved every 2 ps.

The folded structure for the villin headpiece subdomain was obtained from the Protein Data Bank (ID: 1YRF)[S4] and also processed with tleap. For these simulations, frames were saved every ps.

For details on the parameters used in each proposed technique, refer to the Main Text.

#### 2 Supporting Figures

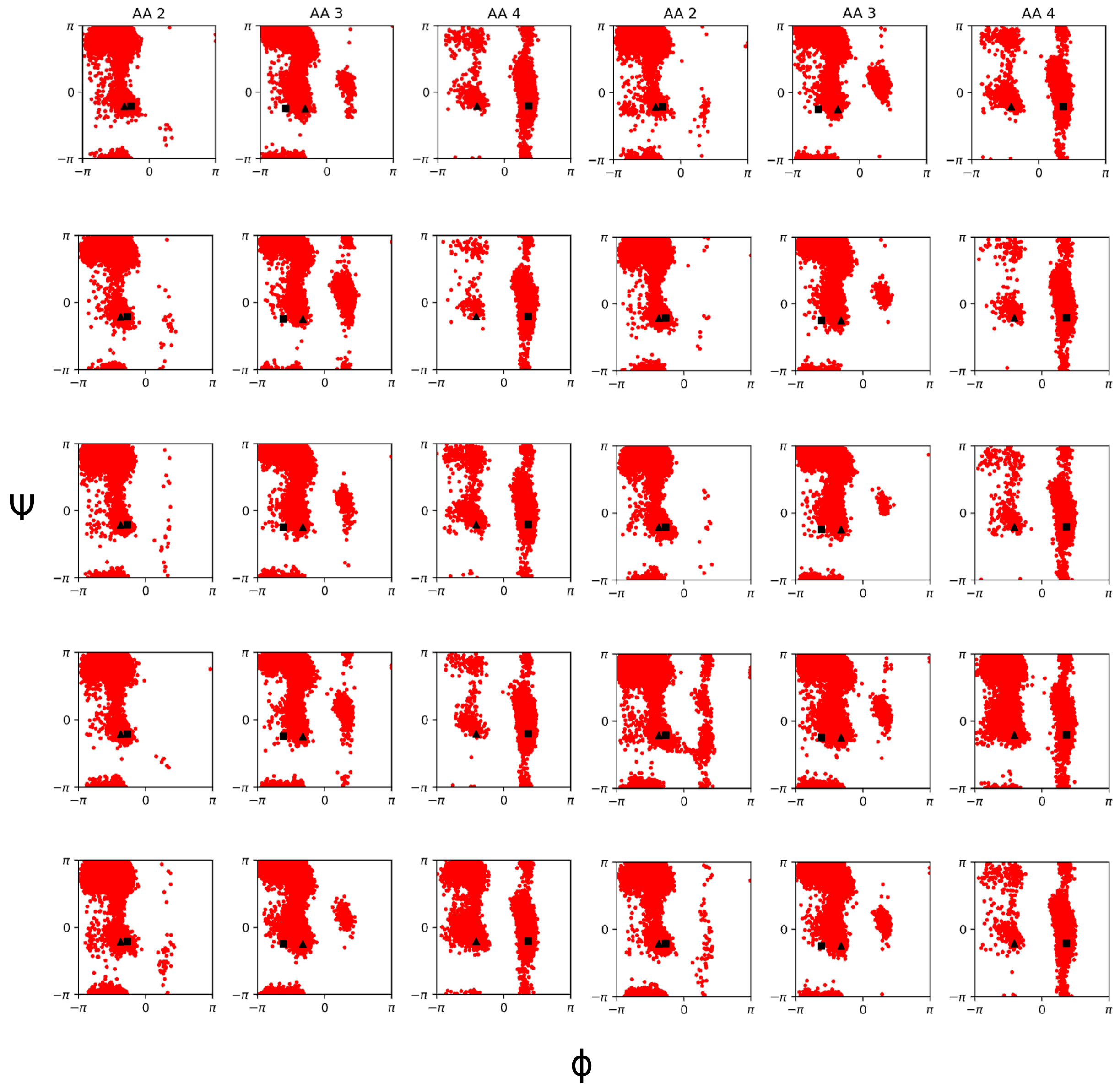


Figure S1: Ramachandran plots for aminoacids L2-A3-L4 in WLALL peptide obtained with Least Counts adaptive sampling (no kinetic model). Replicates 1–10.

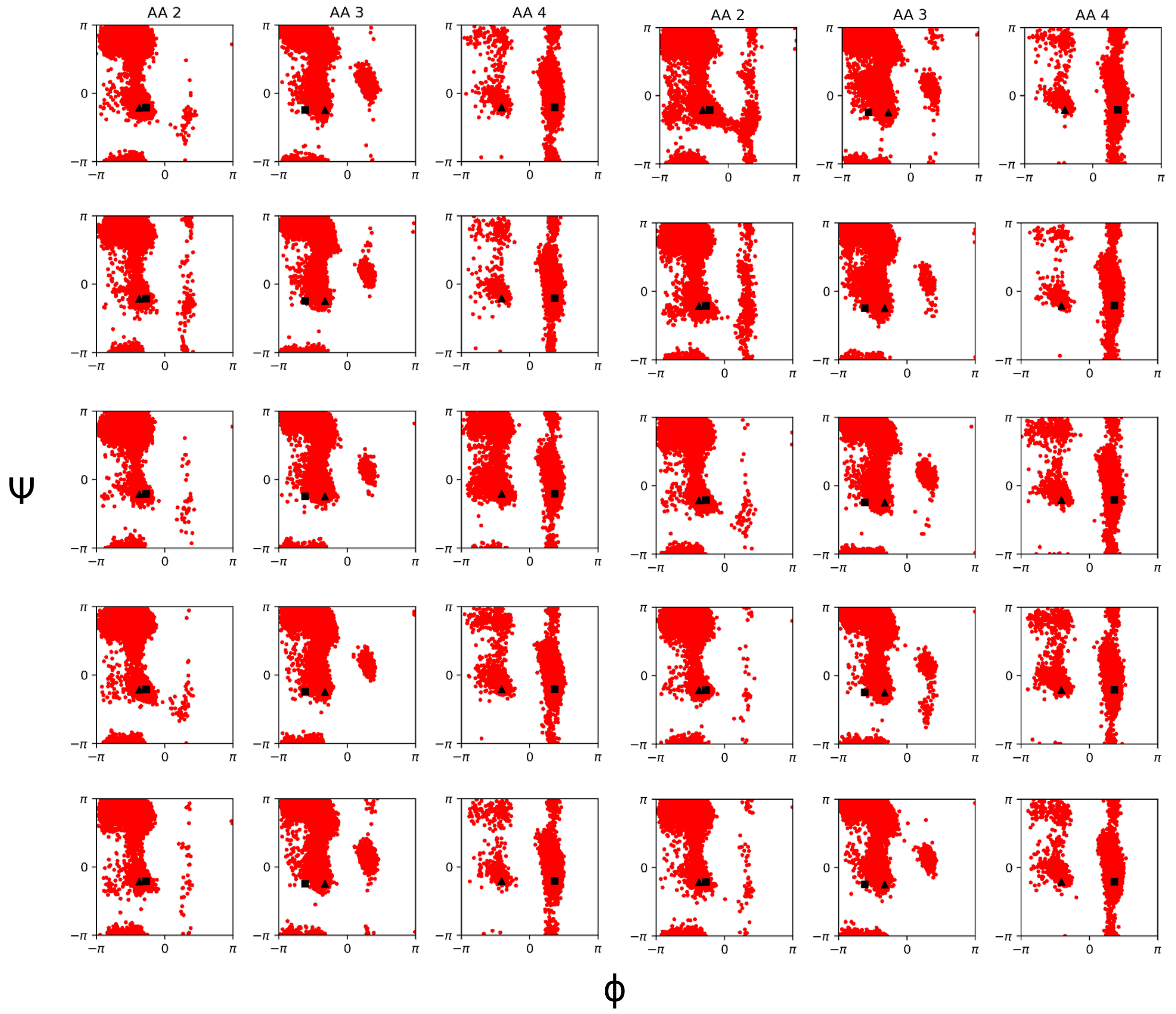


Figure S2: Ramachandran plots for aminoacids L2-A3-L4 in WLALL peptide obtained with Least Counts adaptive sampling (no kinetic model). Replicates 11–20.

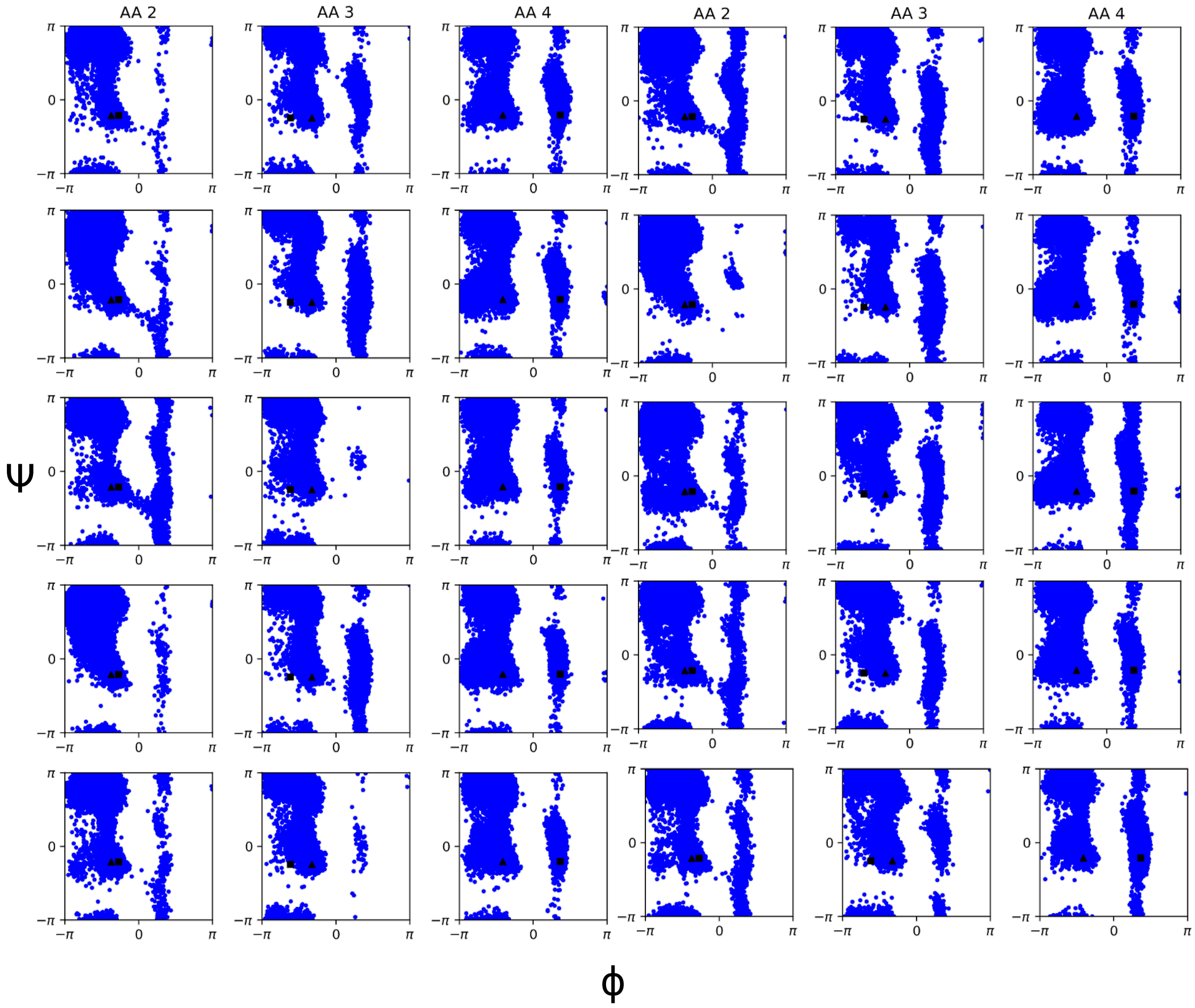


Figure S3: Ramachandran plots for aminoacids L2-A3-L4 in WLALL peptide obtained with VAMPNet + Least Counts (see Section 3.1 in Main Text). Replicates 1–10.

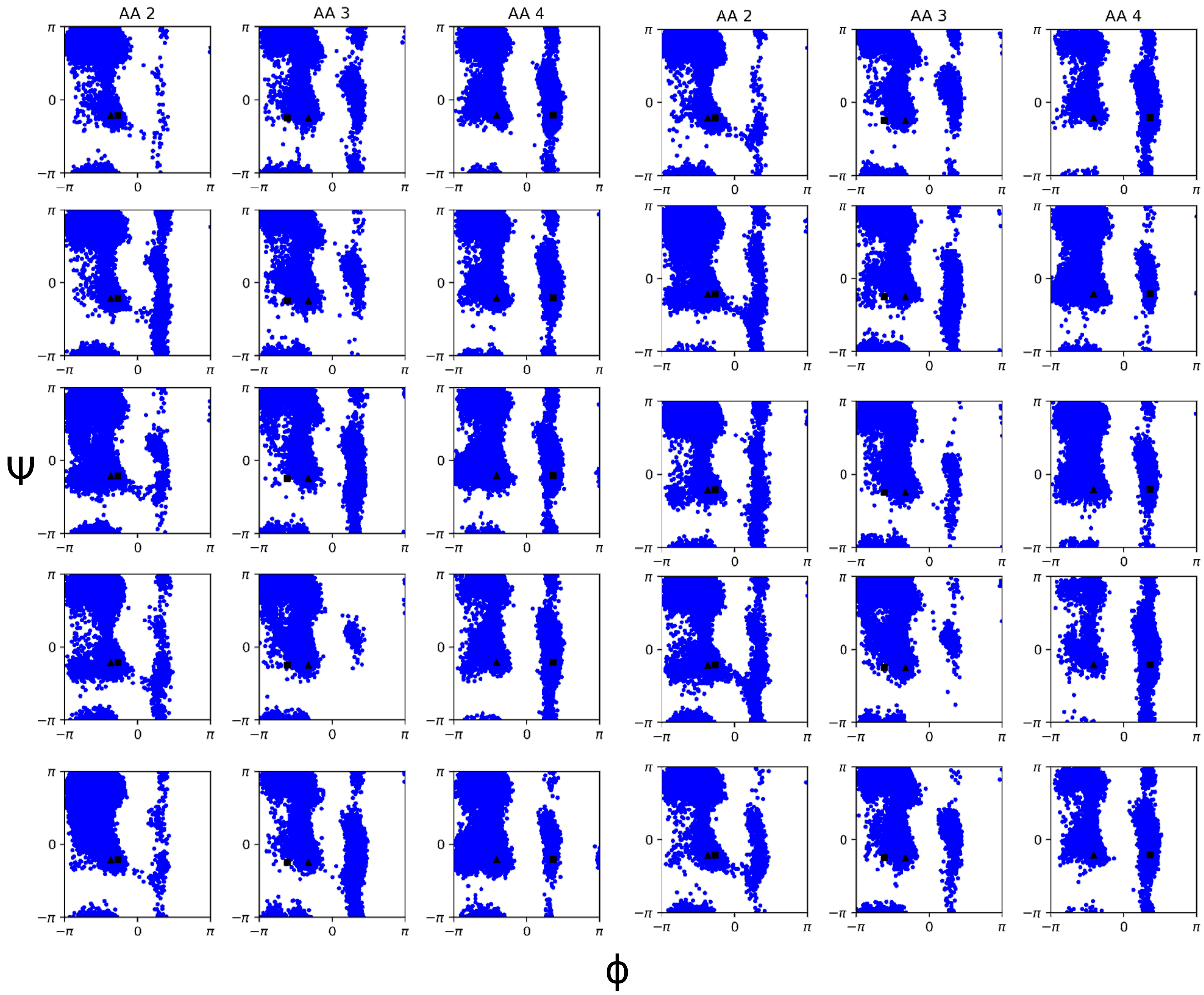


Figure S4: Ramachandran plots for aminoacids L2-A3-L4 in WLALL peptide obtained with VAMPNet + Least Counts (see Section 3.1 in Main Text). Replicates 11–20.

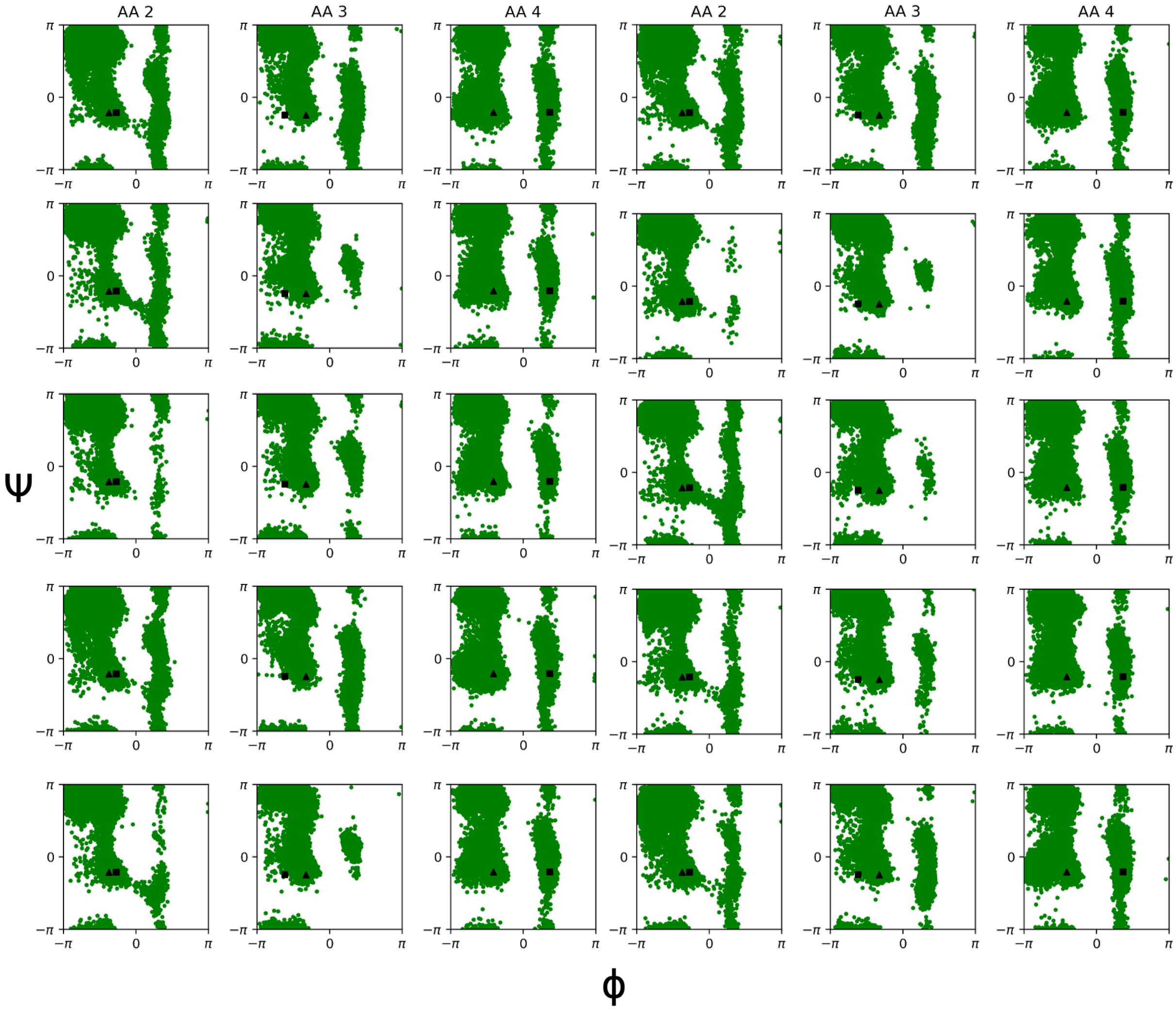


Figure S5: Ramachandran plots for aminoacids L2-A3-L4 in WLALL peptide obtained with VAMPNet + MaxEnt (see Section 3.2 in Main Text). Replicates 1–10.

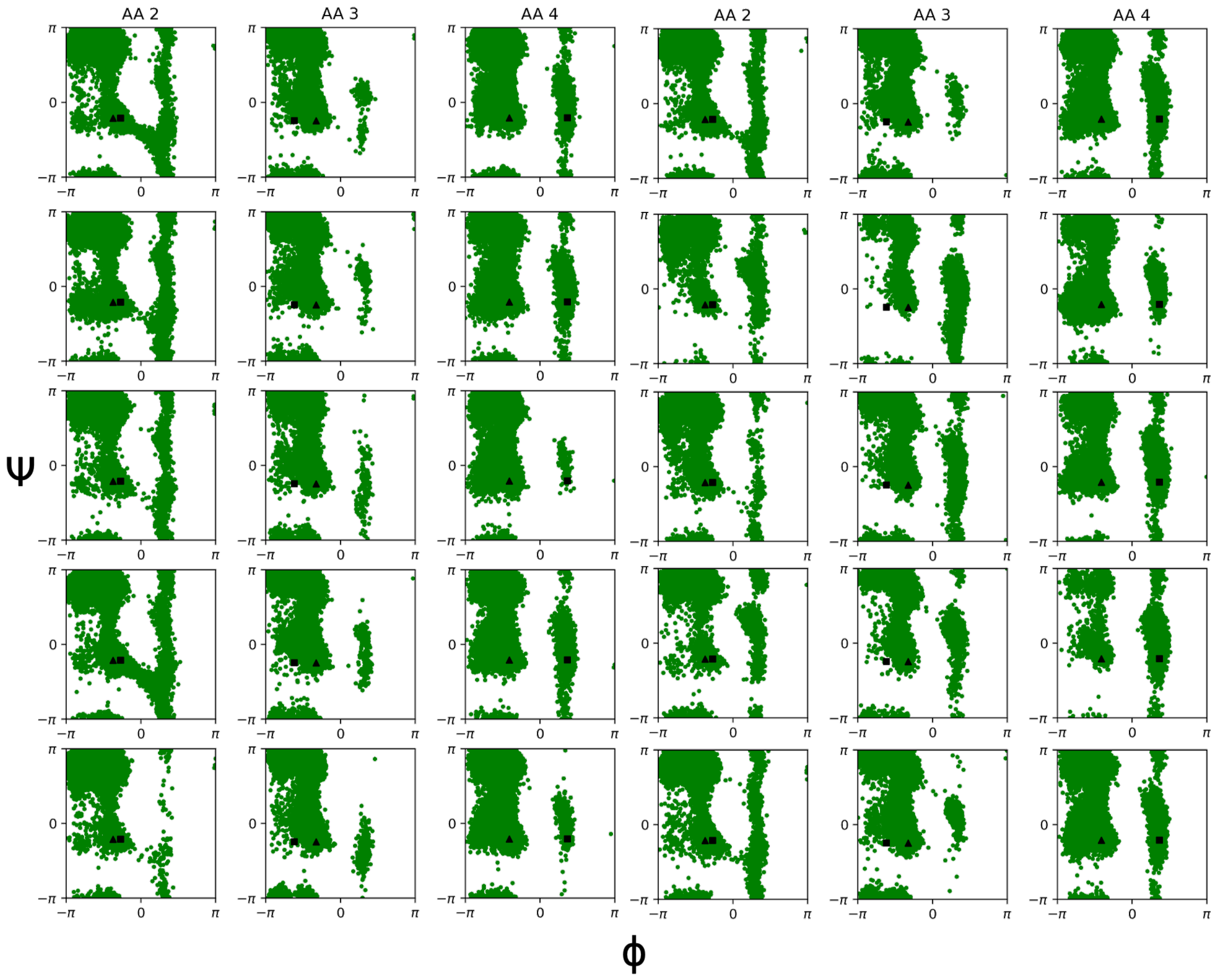


Figure S6: Ramachandran plots for aminoacids L2-A3-L4 in WLALL peptide obtained with VAMPNet + MaxEnt (see Section 3.2 in Main Text). Replicates 11–20.

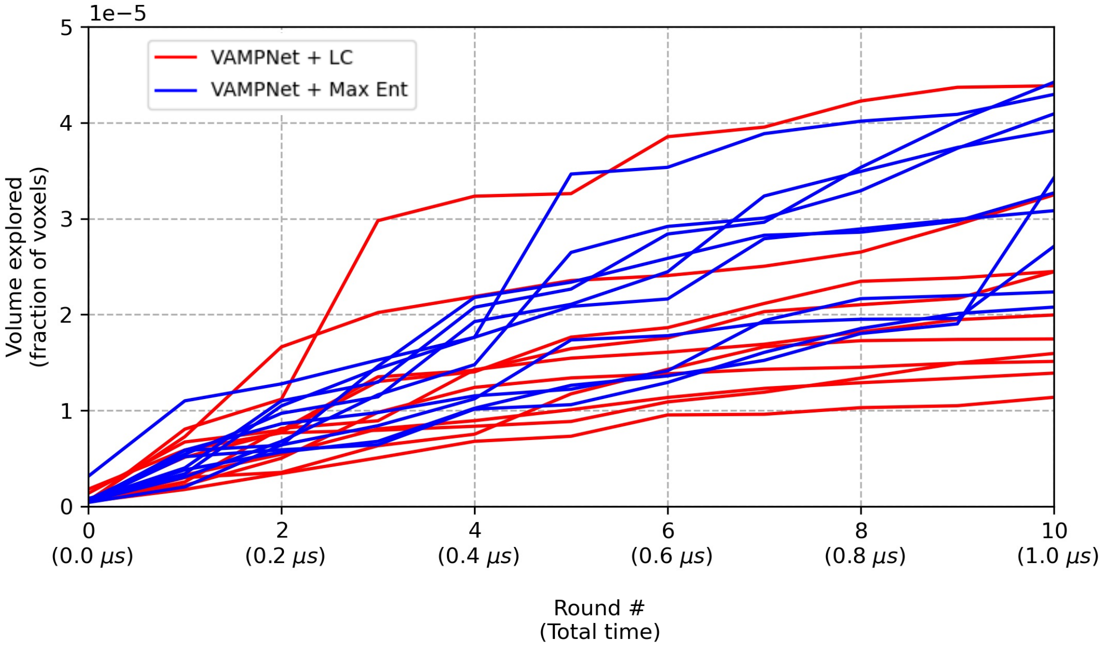


Figure S7: Volume explored in tIC space (as fraction of total voxels) for the villin headpiece subdomain for all replicates of VAMPNet + LC and VAMPNet + MaxEnt.

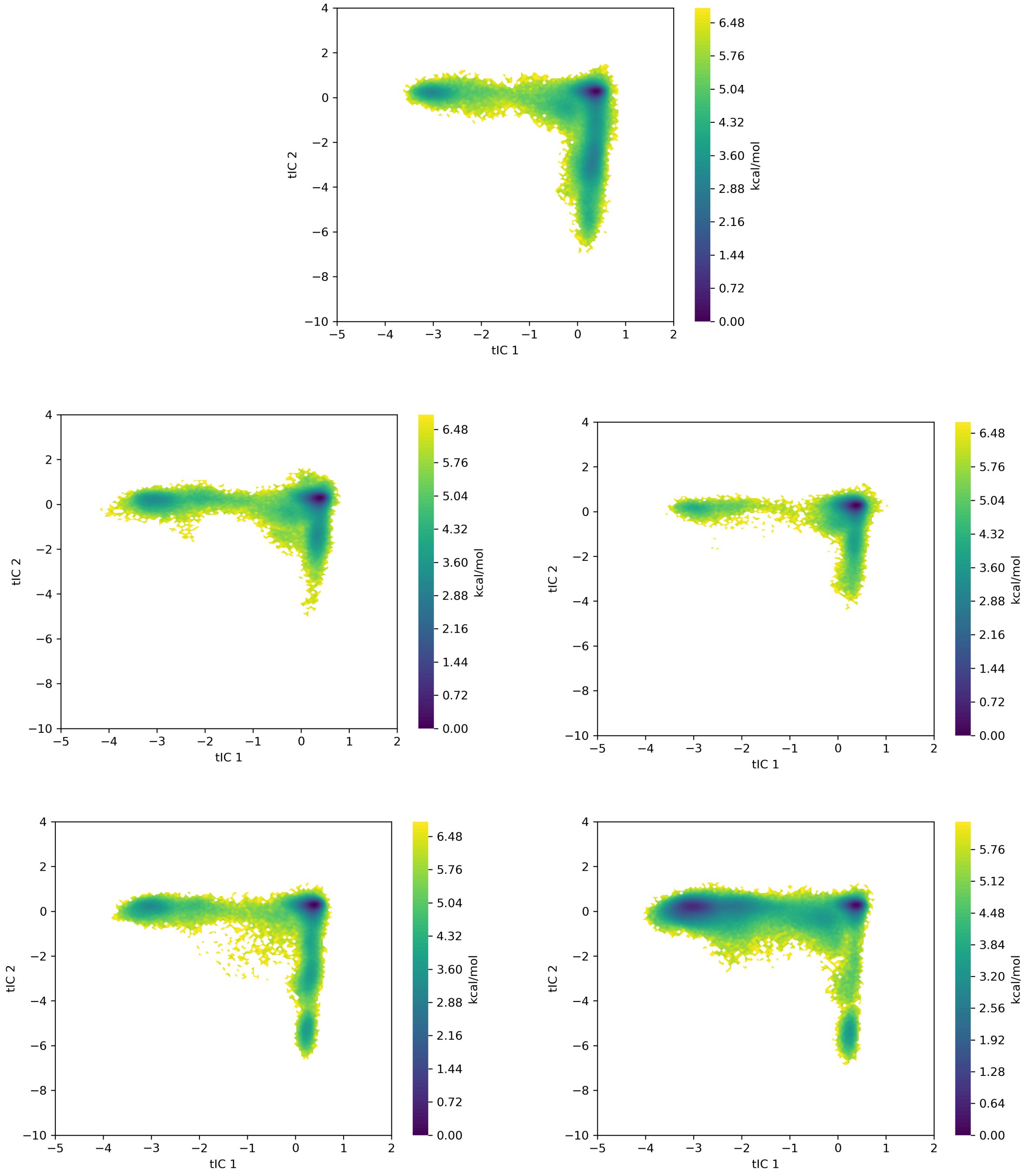


Figure S8: Landscapes in tIC1-tIC2 space for replicates 1–5 of VAMPNet + LC for villin headpiece subdomain.

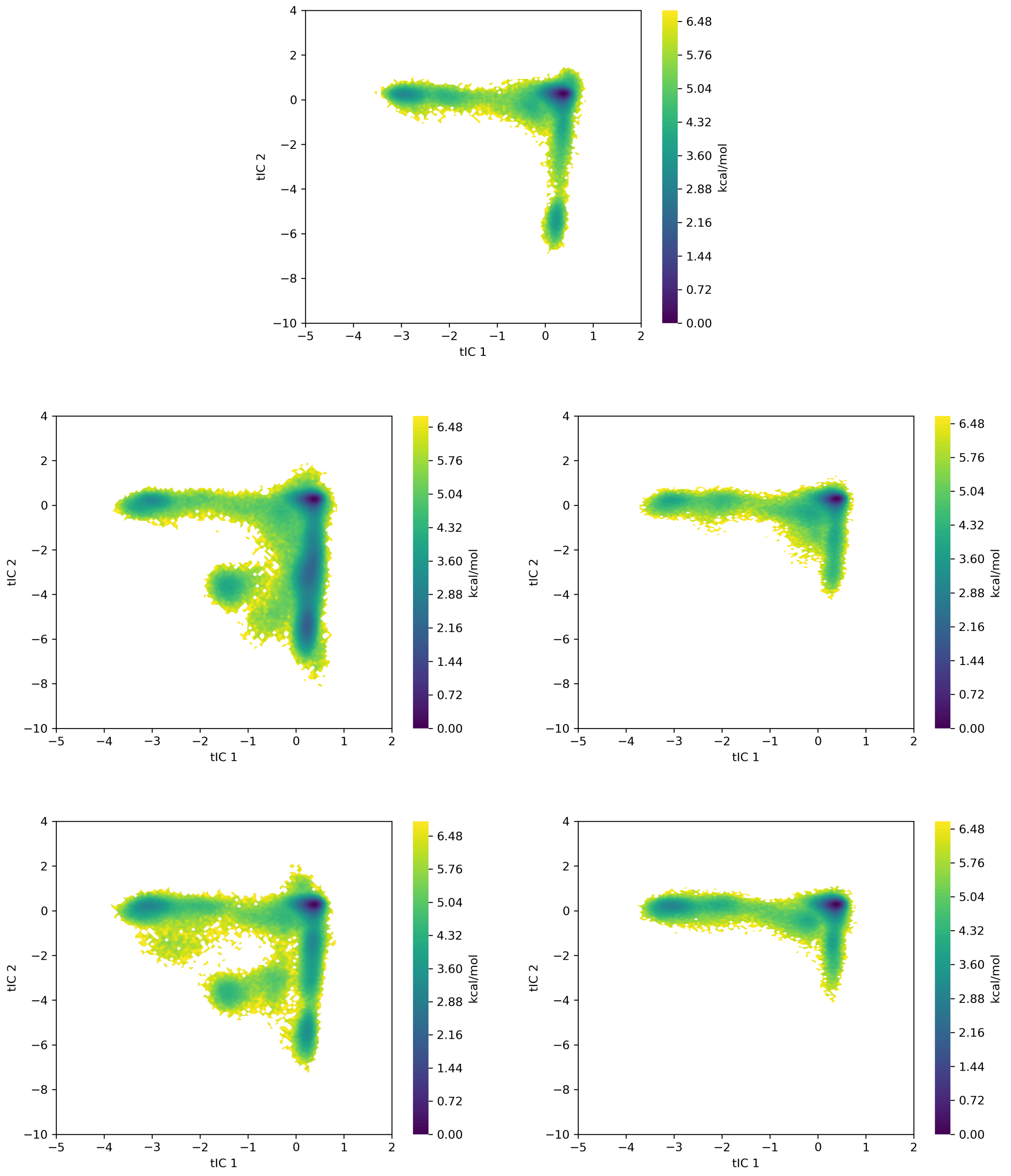


Figure S9: Landscapes in tIC1-tIC2 space for replicates 6–10 of VAMPNet + LC for villin headpiece subdomain.

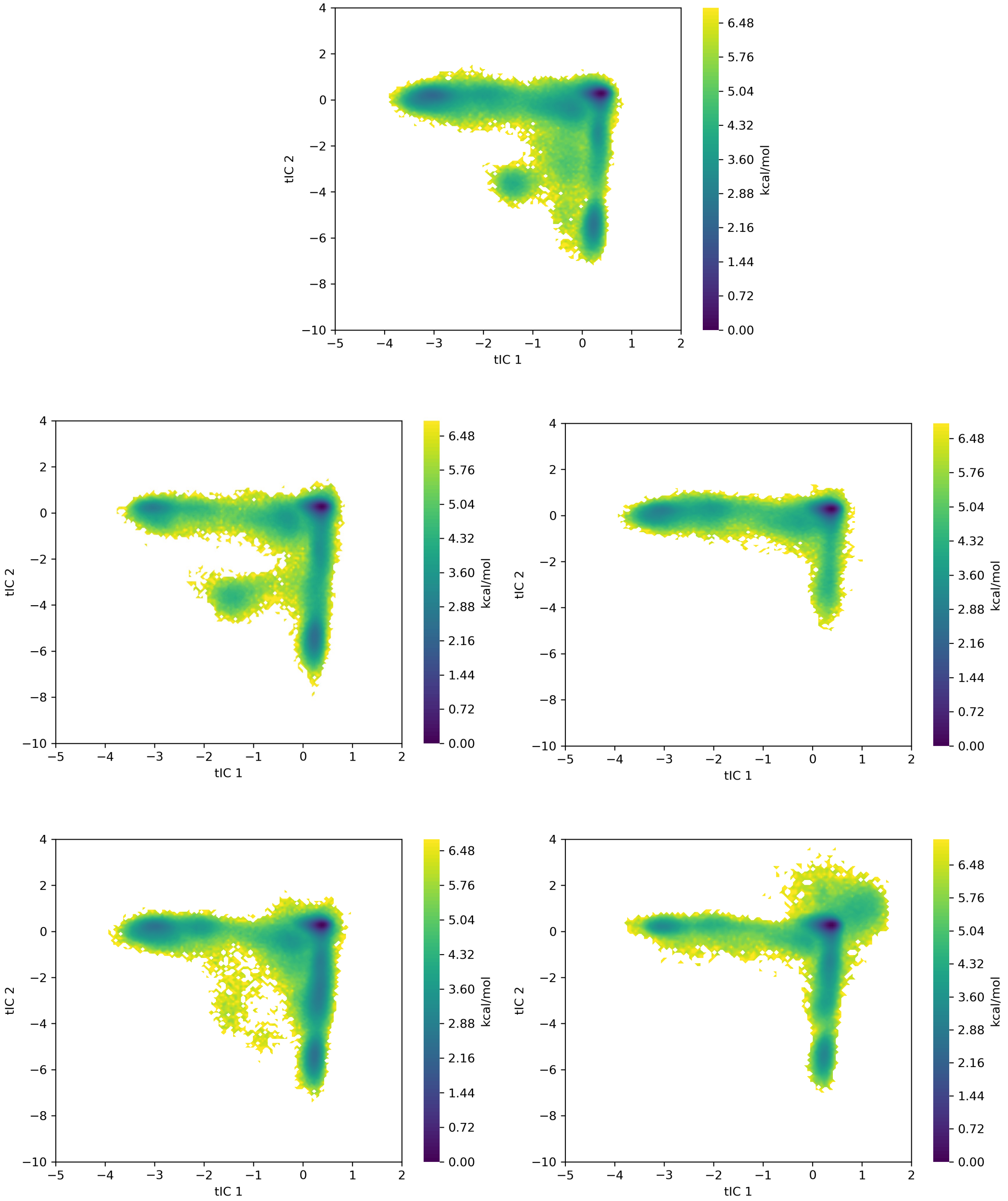


Figure S10: Landscapes in tIC1-tIC2 space for replicates 1–5 of VAMPNet + MaxEnt for villin headpiece subdomain.

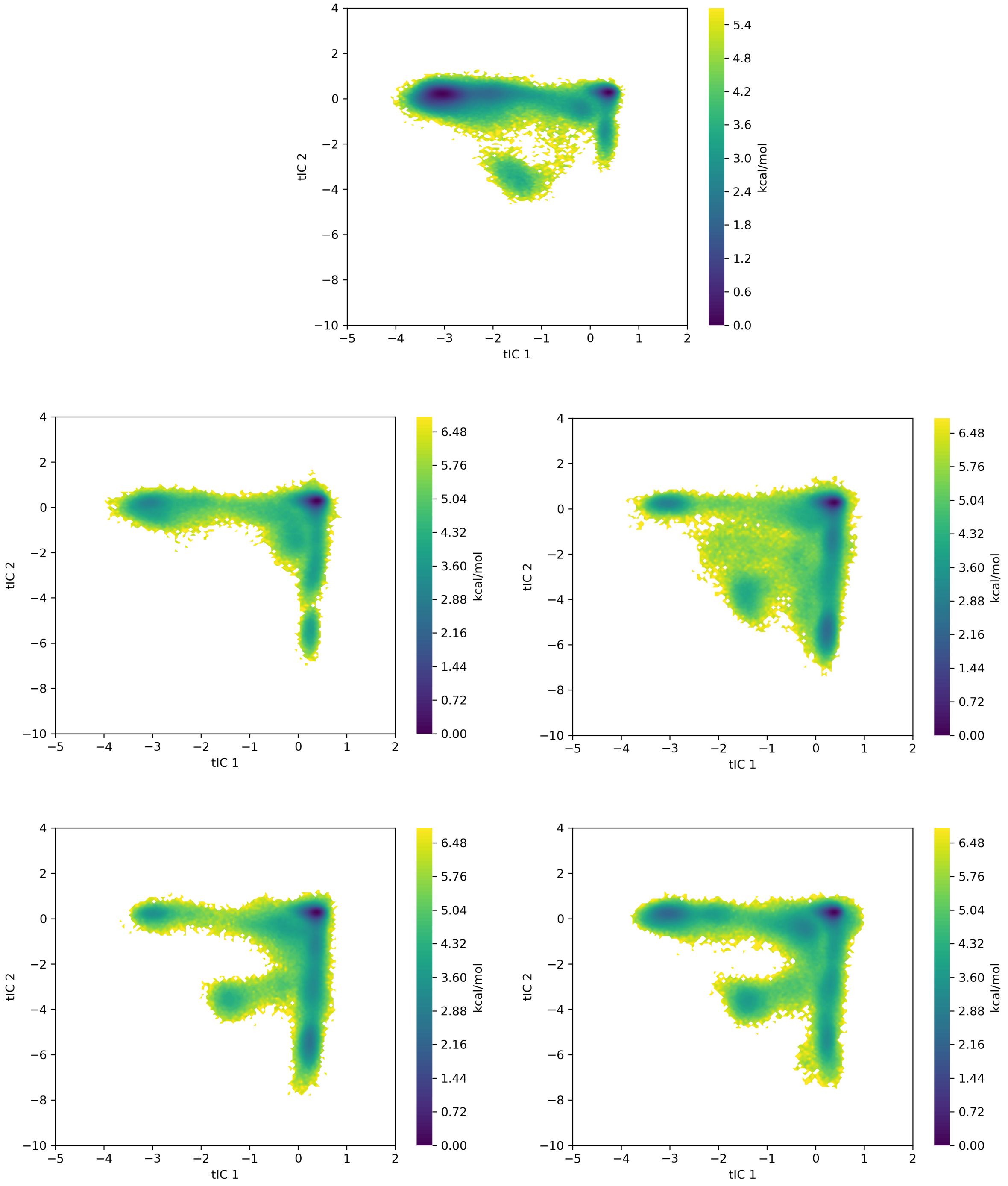


Figure S11: Landscapes in tIC1-tIC2 space for replicates 6–10 of VAMPNet + MaxEnt for villin headpiece subdomain.

### References


(S1) Eastman, P.; Swails, J.; Chodera, J. D.; McGibbon, R. T.; Zhao, Y.; Beauchamp, K. A.; Wang, L.-P.; Simmonett, A. C.; Harrigan, M. P.; Stern, C. D., et al. OpenMM 7: Rapid development of high performance algorithms for molecular dynamics. *PLoS computational biology* **2017**, *13*, e1005659.

(S2) Maier, J. A.; Martinez, C.; Kasavajhala, K.; Wickstrom, L.; Hauser, K. E.; Simmerling, C. ff14SB: improving the accuracy of protein side chain and backbone parameters from ff99SB. *Journal of chemical theory and computation* **2015**, *11*, 3696–3713.

(S3) Scherer, M. K.; Trendelkamp-Schroer, B.; Paul, F.; Pérez-Hernández, G.; Hoffmann, M.; Plattner, N.; Wehmeyer, C.; Prinz, J.-H.; Noé, F. PyEMMA 2: A Software Package for Estimation, Validation, and Analysis of Markov Models. *Journal of Chemical Theory and Computation* **2015**, *11*, 5525–5542.

(S4) Chiu, T. K.; Kubelka, J.; Herbst-Irmer, R.; Eaton, W. A.; Hofrichter, J.; Davies, D. R. High-resolution x-ray crystal structures of the villin headpiece subdomain, an ultrafast folding protein. *Proceedings of the National Academy of Sciences* **2005**, *102*, 7517–7522.
